# Secreted phospholipase A2α generates a pathogen-derived lysophospholipid to signal local immunity

**DOI:** 10.64898/2026.05.15.725072

**Authors:** Hyoung Yool Lee, Tengfang Ling, Jihye Jung, Krishna Kumar, Sang-Chul Kim, Sunyo Jung, Inhwan Hwang, Kyoungwhan Back, Stephen B. Ryu

**Affiliations:** Plant Systems Engineering Research Center, Korea Research Institute of Bioscience and Biotechnology (KRIBB), Daejeon 34141, Republic of Korea; Biosystems and Bioengineering Division, University of Science and Technology (UST), Daejeon 34141, Republic of Korea; R&D Center, Our Bio Ltd. / JICO Ltd., Daejeon 35216, Republic of Korea; School of Life Sciences & Biotechnology, BK21 FOUR KNU Creative BioResearch Group, Kyungpook National University, Daegu 41566, Republic of Korea; Department of Life Sciences, Pohang University of Science and Technology (POSTECH), Pohang 37673, Republic of Korea; Department of Biotechnology, Chonnam National University, Gwangju 61186, Republic of Korea

**Author notes:** These authors contributed equally (Hyoung Yool Lee, Tengfang Ling, Stephen B. Ryu). Division of Agricultural Microbiology, National Institute of Agricultural Science, Rural Development Administration, Wanju 55365, Republic of Korea (Jihye Jung).

**Keywords:** Secreted phospholipase A2α, Pathogen-derived lipid signal, Effector-triggered immunity, Lysophosphatidylethanolamine, Local immunity, *Arabidopsis*

## Abstract

In plants, effector-triggered immunity (ETI) is initiated when NLR receptors recognize pathogen effectors, yet the molecular signals linking this recognition to downstream defense activation remain poorly defined, in contrast to the well-characterized immunogenic signals of pattern-triggered immunity. Here, we show that a lysophospholipid is a previously unidentified ETI-mediating immune signal generated through host enzymatic conversion of pathogen membrane lipids. Upon recognition of avirulent *Pseudomonas syringae*, secretory phospholipase A2α (PLA2α) is rapidly induced and secreted into the apoplast, where it hydrolyzes bacterial phosphatidylethanolamine to produce lysophosphatidylethanolamines (LPEs), predominantly LPE18:1. Genetic ablation of *PLA2α* compromises local immunity, hypersensitive response confinement, and defense gene activation, all of which are largely rescued by exogenous LPE18:1. Mechanistically, LPE18:1 promotes *ICS1*-dependent salicylic acid biosynthesis and *NPR1*-mediated transcriptional reprogramming. Together, these findings support pathogen-derived LPE18:1 as a newly identified lipid-based immune signal that links NLR activation to spatially confined defense responses. The evolutionary conservation of secretory PLA2 enzymes and lysophospholipid signaling from plants to mammals suggests that host-directed enzymatic remodeling of pathogen membranes into immune-activating lipid signals may constitute a conserved strategy of innate immunity.

## INTRODUCTION

Effector-triggered immunity (ETI) in plants is mediated by intracellular nucleotide-binding leucine-rich repeat (NLR) receptors, which recognize pathogen-derived effectors and activate defense responses. When *Pseudomonas syringae* pv. *tomato* DC3000 carrying AvrRpm1 (*Pst AvrRpm1*) infects the extracellular spaces of host plants, the bacterium delivers the AvrRpm1 protein into the host cytoplasm via the type III secretion system (Chisholm et al., 2006; Wang et al., 2022). If the host plant expresses the NLR receptor RPM1, it indirectly perceives AvrRpm1 and initiates defense responses. NLR–effector recognition triggers early immune signaling events, including conformational activation of NLRs into higher-order resistosome complexes, calcium ion (Ca²⁺) influx, reactive oxygen species (ROS) production, and transcriptional reprogramming leading to ETI (Mammarella et al., 2015; Zhou and Zhang, 2020; Dodds et al., 2024; Jones et al., 2024).

The local defense response induced by ETI includes the hypersensitive response (HR), which involves localized programmed cell death at the infection site (Fu and Dong, 2013; Zhou et al., 2023; Dodds et al., 2024). The HR actively restricts biotrophic pathogen proliferation by depriving them of nutrients and confining infections. HR is tightly regulated by immune signaling networks, including NLR receptor activation and downstream executor mechanisms (Contreras et al., 2023). Several studies underscore the importance of maintaining ROS homeostasis for proper HR execution (Kaur et al., 2021; Ngou et al., 2021; Ugalde, 2023). Thus, HR represents a genetically encoded and dynamically regulated defense strategy central to plant local resistance.

Salicylic acid (SA) serves as a key regulator of ETI and HR. The plastidial isochorismate synthase 1 (ICS1) pathway represents the major source of SA biosynthesis in *Arabidopsis* during avirulent *Pseudomonas* infection. Mutants defective in *ICS1* (*sid2*) fail to accumulate SA and exhibit strongly compromised HR and pathogen resistance (Nawrath and Metraux, 1999; Dewdney et al., 2000; Zhang et al., 2010; Li et al., 2023). Moreover, the master regulator NPR1 is essential for SA-dependent transcriptional activation of *PATHOGENESIS-RELATED* (*PR*) genes and for restricting pathogen growth (Tada et al., 2008; Liu et al., 2020; Dong, 2025; Zhang et al., 2025). Since SA biosynthesis and NPR1 activation are essential for robust defense outputs and transcriptional reprogramming, elucidating the upstream signals that mediate these processes during ETI remains a top research priority.

The apoplast represents a critical battlefield between plants and pathogens, serving not only as a physical barrier but also as a site where secreted molecules contribute to immune responses. These include hydrolytic enzymes (e.g., glucanases and chitinases) and proteases that degrade pathogen-derived molecules, which are governed by secretory pathways (Wan et al., 2021; Del Corpo et al., 2024). Growing evidence indicates that secretory proteins and trafficking pathways play pivotal roles in plant immunity (Yun et al., 2023; Bhandari and Brandizzi, 2024). Notably, the *P. syringae* effector AvrRpm1 suppresses plant immune secretion by interfering with the exocyst complex, thereby reducing the secretion of defense molecules such as callose (Redditt et al., 2019). Based on these previous findings, it is plausible that ETI involves specific secretory proteins that contribute to local immune responses. However, the molecular mechanisms linking the secretory pathway to ETI signaling remain largely unknown.

Lipid signaling has emerged as a potential regulatory module in ETI, but its role remains poorly understood (Shah, 2005; Lim et al., 2017; Vӧlz et al., 2021; Song et al., 2022; Yang et al., 2022). For instance, lysophospholipids such as lysophosphatidylcholine (LPC) have been shown to mediate plant symbiosis with arbuscular mycorrhizal fungi (Drissner et al., 2007; Vijayakumar et al., 2016). Furthermore, studies in animal systems have demonstrated that lysophospholipids, such as lysophosphatidic acid (LPA), function as signaling molecules, facilitating cell-to-cell communication to activate immune responses and developmental processes (Tan et al., 2020; Butera et al., 2022). These observations suggest that lysophospholipids may function in plant immune signaling. Our earlier work revealed that lysophosphatidylethanolamine (LPE), a lysophospholipid generated by phospholipase A2 (PLA2), induces *PR* gene expression and enhances disease resistance against both bacterial and fungal pathogens when applied exogenously to plant leaves (US patent 6,559,099, 2003; US patent 10,030,256, 2018). A secretory PLA2 isoform, PLA2α, is translocated from Golgi bodies to the apoplast as leaves mature, a process that is further accelerated by bacterial infection (Jung et al., 2012). Similar protective effects of exogenously applied LPE have independently been reported (Vӧlz et al., 2021; Vӧlz et al., 2023). However, a direct mechanistic link between PLA2 enzymes and their lipid-derived immune functions has not yet been fully established.

Here, we establish that secretory PLA2α and its lipid product, LPE, serve as essential mediators of local immune responses in *Arabidopsis*. Our findings address this gap by establishing LPE18:1 as a novel ETI-mediating immune signal—generated not from pre-existing pathogen-associated molecular patterns, but through the active enzymatic conversion of pathogen membrane components by the host phospholipase PLA2α. Mechanistically, we propose that apoplastic LPE18:1 acts as a lipid signal that reinforces local defense activation at and around infection sites, thereby establishing LPE18:1 as a previously unrecognized molecular link between NLR-mediated effector recognition and the execution of spatially confined ETI responses.

## RESULTS

### PLA2α is rapidly expressed and translocates to the apoplast upon *Pst AvrRpm1* inoculation

The apoplast represents a primary interface of plant–pathogen interactions, positioning apoplastic lipid-metabolizing enzymes such as secretory phospholipase A2s as key candidates for early immune regulation. Among the four secretory PLA2 isoforms in *Arabidopsis* (Ryu, 2004), only *PLA2α* was rapidly induced 1–3 hours post-inoculation (hpi) with *Pst AvrRpm1* (Figures 1A and S1A). Subcellular localization assays using transgenic plants co-expressing PLA2α-RFP and the Golgi marker ST-GFP revealed enhanced translocation of PLA2α to the apoplast upon inoculation with *Pst AvrRpm1*. This localization pattern was further verified by cell wall staining with FB-28 after plasmolysis (Figures 1B and S1B), consistent with our previous findings (Jung et al., 2012). Immunoblot analysis revealed that PLA2α was enriched approximately two-fold in the apoplastic fractions of *Pst AvrRpm1*-inoculated leaves compared to mock-treated controls (Figures 1C and S1C). The malate dehydrogenase (MDH) assay confirmed no cytosolic contamination in the apoplastic fractions (Figure S1D). Together, these results indicate enrichment of PLA2α in the apoplasts of *Pst AvrRpm1-*infected leaves.

**Figure 1.**
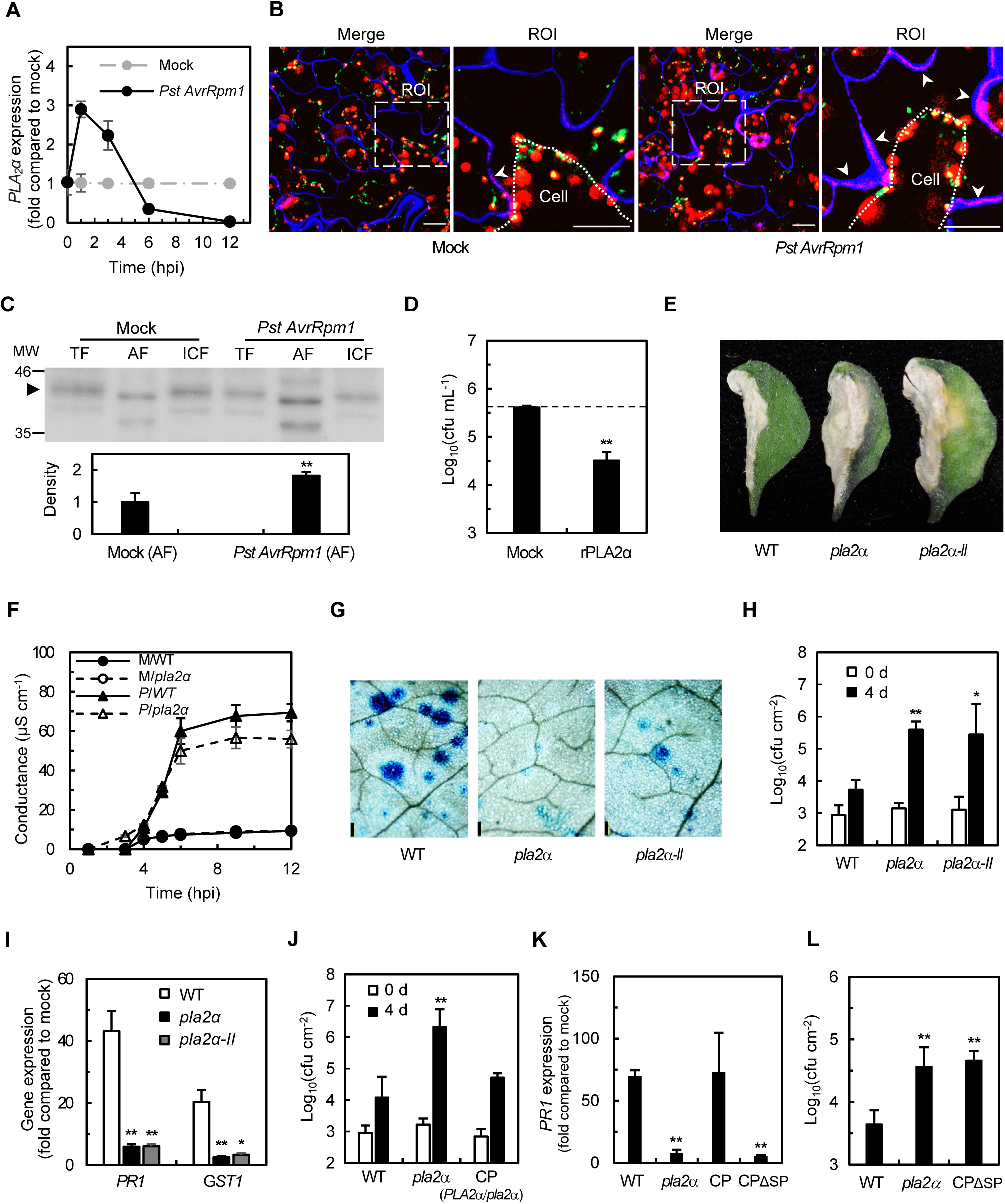
PLA2α is induced and secreted during ETI and contributes to local immunity. **(A)** Time-course analysis of *PLA2α* expression during ETI. Wild-type plants were spray-inoculated with *Pst AvrRpm1* (1 × 10^8^ CFU mL⁻¹), and *PLA2α* expression was quantified by RT-qPCR at the indicated time points. Data are expressed as fold-change relative to mock-inoculated plants. **(B)** Subcellular localization of PLA2α-DsRed2 in leaf cells before and after inoculation with *Pst AvrRpm1*. Transgenic plants co-expressing PLA2α-DsRed2 (red) and ST-GFP (Golgi marker, green) were spray-inoculated with *Pst AvrRpm1* (1 × 10^8^ CFU mL⁻¹). At 3 hpi, leaves were infiltrated with FB-28 (0.1 mg mL⁻¹) for cell wall staining (blue) and plasmolyzed in 1 M KNO_3_ prior to imaging. "Merge" shows the overlay of three-channel images; "ROI" is a magnification of the dashed rectangle in "Merge." Dashed lines in "ROI" mark the plasmolyzed plasma membrane. Arrowheads indicate co-localization of PLA2α-DsRed2 with the cell wall in the apoplast. Scale bar, 20 μm. **(C)** Immunoblot analysis of PLA2α-DsRed2 in subcellular fractions from leaves in (B), detected with an anti-DsRed2 antibody. TF, total fraction; AF, apoplastic fraction; ICF, intracellular fraction. The arrowhead indicates the expected molecular weight of PLA2α-DsRed2 (42.1 kDa). Band intensities in AF relative to mock are indicated below the blot. **(D)** Antimicrobial activity of recombinant PLA2α (rPLA2α) against *Pst*. Bacterial suspensions (5 × 10⁵ CFU mL⁻¹) were incubated with purified rPLA2α or mock for 6 h, and surviving bacteria were quantified by CFU assay. The dashed line indicates the CFU count at 0 h. **(E)** MacroHR and disease symptoms in WT and *pla2α* mutants following *Pst AvrRpm1* infection. Leaves were syringe-infiltrated with *Pst AvrRpm1* (5 × 10⁷ CFU mL⁻¹) and photographed at 6 dpi. **(F)** Time-course analysis of electrolyte leakage in WT and *pla2α* during *Pst AvrRpm1*-triggered ETI. Leaf discs were vacuum-infiltrated with *Pst AvrRpm1* (5 × 10⁷ CFU mL⁻¹) or mock, and conductance was measured at the indicated time points up to 12 hpi. M, mock-inoculated; P, *Pst AvrRpm1*-inoculated. **(G)** Trypan blue staining of microHR cell death in WT and *pla2α* mutants. Plants were spray-inoculated with *Pst AvrRpm1* (4 × 10⁸ CFU mL⁻¹), and leaves were stained with trypan blue at 12 hpi and visualized by light microscopy. Scale bar, 200 μm. **(H)** Bacterial growth in WT and *pla2α* mutants following *Pst AvrRpm1* spray inoculation. Plants were spray-inoculated with *Pst AvrRpm1* (4 × 10⁸ CFU mL⁻¹), and bacterial populations in leaf discs were quantified by CFU assay at the indicated dpi. **(I)** Expression of defense marker genes *PR1* and *GST1* in WT and *pla2α* mutants following *Pst AvrRpm1* infection. Leaves were syringe-infiltrated with *Pst AvrRpm1* (1 × 10⁷ CFU mL⁻¹), and gene expression was quantified by RT-qPCR at 8 hpi. Data are expressed as fold-change relative to mock-infiltrated plants. **(J)** Bacterial growth in *pla2α* complemented with native PLA2α (CP). Plants were spray-inoculated with *Pst AvrRpm1* (4 × 10⁸ CFU mL⁻¹), and bacterial populations were quantified by CFU assay at the indicated dpi. **(K)** *PR1* expression in *pla2α* complemented with full-length PLA2α (CP) or signal-peptide-deleted PLA2α (CPΔSP). Leaves were syringe-infiltrated with *Pst AvrRpm1* (1 × 10⁷ CFU mL⁻¹), and *PR1* expression was quantified by RT-qPCR at 8 hpi. Data are expressed as fold-change relative to mock-infiltrated plants. **(L)** Bacterial growth in CPΔSP plants at 4 dpi. Plants were spray-inoculated with *Pst AvrRpm1* (4 × 10⁸ CFU mL⁻¹), and bacterial populations were quantified by CFU assay at 4 dpi. Data represent means ± SD (A, C, D, F, H, J, and L) or ± SEM (I and K) from at least three independent experiments. Asterisks indicate statistically significant differences from WT by two-tailed Student’s *t*-test (\**P* < 0.05; \*\**P* < 0.01). See also Figures S1 and S2.

Consistent with the known ability of PLA2α enzyme to hydrolyze membrane phospholipids (Ryu et al., 2005), recombinant PLA2α (rPLA2α) protein exhibited strong antimicrobial activity in an *in vitro* colony-forming units (CFU) assay (Figure 1D). After 6 h of incubation, rPLA2α reduced bacterial viability to ∼6% of the initial count, while mock treatment had no detectable effect, indicating that rPLA2α exhibits direct antimicrobial activity against *Pst AvrRpm1 in vitro*.

### Effector-triggered immune responses are impaired in *pla2α*

Given that *Pst AvrRpm1* colonizes the apoplastic space, it is plausible that PLA2α engages in dynamic molecular interactions with invading pathogens in this compartment, where host–pathogen exchanges are highly active. To investigate the cellular function of secretory PLA2α in plant defense, we examined *Arabidopsis pla2α* (knockout) and *pla2α-II* (knockdown) mutants obtained from TAIR (Figures S2A and S2B) in response to pathogen infection. Under normal growth conditions, the *pla2α* mutants showed no detectable differences in phenotype compared to Col-0 wild-type (WT) plants. However, upon infection with *Pst AvrRpm1*, the *pla2α* mutants exhibited markedly different phenotypes. When half-leaves of WT, *pla2α*, and *pla2α-II* plants were infiltrated with a bacterial dose (5 × 10⁷ CFU mL^-1^), although the initial phenotypic HR appeared almost normal in the mutants compared to WT (Figure S2C), the *pla2α* and *pla2α-II* lines failed to form distinct boundaries around the HR region. Instead, lesions expanded beyond the initially infiltrated region in *pla2α* mutants, accompanied by yellowing of the surrounding leaf tissue, whereas WT plants maintained sharply confined HR boundaries (Figure 1E and S2D).

To further assess HR, ion leakage assays were performed using two different inoculation methods. In the detached leaf disc assay with vacuum-infiltration, HR induced by *Pst AvrRpm1* was attenuated at early stages of cell death in *pla2α* compared to WT plants. Similar reductions were observed following infiltrations with *Pst AvrRpt2* and *Pst AvrRps4* (Figures 1F and S2E). By contrast, in intact leaves syringe-infiltrated with *Pst AvrRpm1*, while WT leaves showed decreased leakage after 1 day post-inoculation (dpi), *pla2α* mutants exhibited a continuous increase up to 4 dpi (Figure S2F). This late increase likely reflects cumulative membrane damage resulting from failure to confine HR-associated cell death, leading to lesion expansion beyond the initially infected region, with accompanying yellowing in surrounding tissues (Figure 1E). Together, these results show that vacuum-infiltrated leaf discs, which experience minimal mechanical stress, sensitively detect early HR defects, whereas high-dose syringe infiltration of intact leaves can mask early differences but clearly reveals late-stage tissue collapse and boundary failure. Thus, while each assay has its advantages, gentler inoculation methods are more suitable for probing early local immune responses.

To investigate the impairment of *pla2α* mutants in local immune responses under more physiologically relevant conditions, we assessed HR and disease symptoms following spray inoculation of leaf surfaces with *Pst AvrRpm1*, which mimics the natural stomatal entry route of the pathogen. Trypan blue staining at 12 hpi revealed that mutant leaves exhibited a reduced number of microHR lesions compared to WT leaves, further supporting a partial defect in HR execution (Figure 1G and S2G). Moreover, visible disease symptoms developed in older leaves of the mutant plants, whereas WT plants exhibited little to no lesions (Figure S2H), indicating a compromised defense phenotype in the absence of functional PLA2α. Consistently, bacterial growth assays revealed significantly higher titers of *Pst AvrRpm1* in both *pla2α* and *pla2α-II* plants compared to WT (Figure 1H), with similar trends observed for *Pst AvrRpt2* and *Pst AvrRps4* (Figure S2I). These results together indicate that *pla2α* mutants exhibit impaired defense responses against bacterial effectors recognized by both CC-NB-LRR and TIR-NB-LRR classes of NLRs.

Supporting this, *pla2α* and *pla2α-II* mutants exhibited significantly reduced expression of defense genes *PR1* and *GST1* after *Pst AvrRpm1* infection compared to WT (Figure 1I). However, when the *pla2α* mutant was complemented with a native promoter-driven pPLA2α*:PLA2α* construct, its ability to induce defense genes and to restrict excessive bacterial growth was restored to levels comparable to those observed in the WT (Figures 1J and 1K). A similar phenotype has been reported for *ICS1* mutant *sid2*, which exhibit partially impaired HR in response to avirulent *Pst* and fail to adequately restrict bacterial growth (Nawrath and Metraux, 1999; Al-Daoude et al., 2005).

To validate the requirement for apoplastic translocation of PLA2α in ETI, a complementation construct (*pPLA2α:PLA2α*ΔSP) was generated in which the signal peptide sequence was deleted, preventing entry into the secretory pathway and retaining the truncated protein in the cytosol. Unlike the full-length complementation construct (CP), CPΔSP failed to restore *PR1* expression or bacterial resistance in *pla2α* (Figures 1K and 1L), demonstrating that apoplastic secretion of PLA2α is essential for its immune function.

### Expression of *PLA2α* is followed by an increase in LPE levels

As shown in Figure 1A, *PLA2α* expression was rapidly induced 1–3 hpi with *Pst AvrRpm1*, whereas virulent *Pst* caused only a slight, non-significant increase at 1 h compared to mock treatment (Figure 2A). Histochemical β-glucuronidase (GUS) assays revealed that *PLA2α* is preferentially expressed at sites infiltrated with *Pst AvrRpm1*, but not at mock- or virulent *Pst*-treated sites, 1.5 hpi, in pPLA2α*:GUS* transgenic *Arabidopsis* leaves (Figure 2B). Together, these findings indicate that *PLA2α* expression is tightly regulated by ETI signaling and is spatially confined to pathogen infection sites.

**Figure 2.**
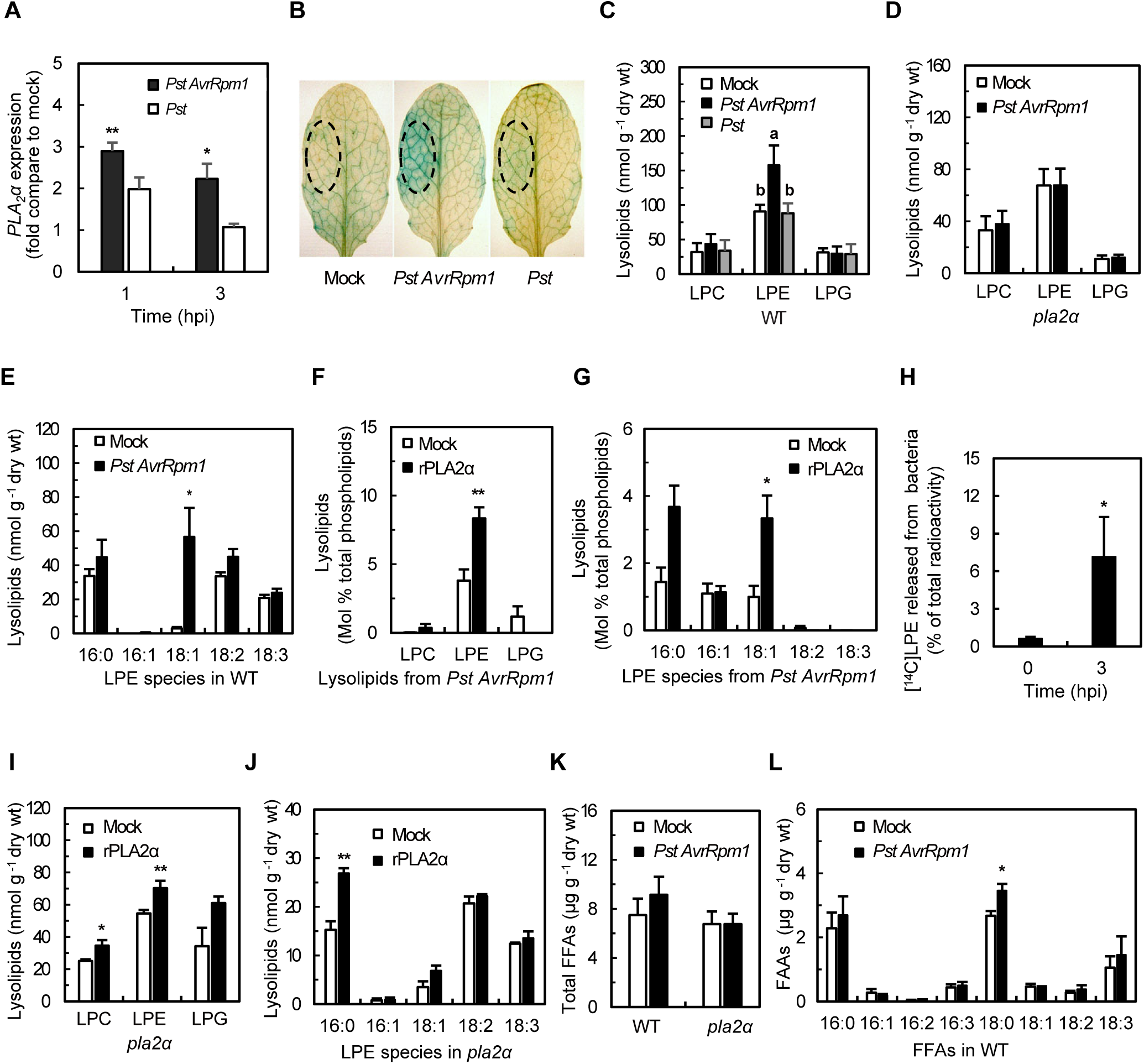
LPE18:1 is generated from invading bacteria by host PLA2α during ETI. **(A)** Expression levels of *PLA2α* in WT plants infected with virulent *Pst* or *Pst AvrRpm1*. Plants were spray-inoculated with *Pst* or *Pst AvrRpm1* (1 × 10⁸ CFU mL⁻¹), and *PLA2α* expression was quantified by RT-qPCR at the indicated hpi. Data are expressed as fold-change relative to mock-inoculated plants. **(B)** Spatial pattern of *PLA2α* induction revealed by GUS reporter assay. Leaves of transgenic plants expressing *PLA2α*::GUS were syringe-infiltrated with *Pst* or *Pst AvrRpm1* (1 × 10⁷ CFU mL⁻¹) and stained at 1.5 hpi. Dashed circles indicate infiltrated areas. **(C–E)** Lysophospholipid profiles in *Pst*- or *Pst AvrRpm1*-infected WT and *pla2α* plants. Plants were spray-inoculated with *Pst* or *Pst AvrRpm1* (1 × 10⁸ CFU mL⁻¹). At 3 hpi, total lipids were extracted and quantified by ESI-MS/MS for total lysophospholipid classes in WT (C) and *pla2α* (D), and for LPE molecular species in WT (E). **(F and G)** Lysophospholipid generation from *Pst AvrRpm1* lipids by rPLA2α *in vitro*. Total lipids extracted from *Pst AvrRpm1* were incubated with rPLA2α for 15 min and quantified by ESI-MS/MS for lysophospholipid classes (F) and LPE molecular species (G). Values are expressed as mol% of total phospholipids. **(H)** Generation of [¹⁴C]LPE from *Pst AvrRpm1* membrane phospholipids in planta. Leaves were syringe-infiltrated with [¹⁴C]ethanolamine-labeled *Pst AvrRpm1* (1 × 10⁷ CFU mL⁻¹). At the indicated hpi, total lipids were extracted, separated by TLC, and [¹⁴C]LPE radioactivity was quantified by scintillation counting. **(I and J)** Lysophospholipid accumulation in *pla2α* leaves treated with rPLA2α. Leaves were syringe-infiltrated with rPLA2α (10 μg mL⁻¹), and total lipids were extracted after 3 h and quantified by ESI-MS/MS for lysophospholipid classes (I) and LPE molecular species (J). **(K and L)** Free fatty acid (FFA) levels in WT and *pla2α* following *Pst AvrRpm1* infection. Plants were spray-inoculated with *Pst AvrRpm1* (1 × 10⁸ CFU mL⁻¹), and total FFAs (K) and FFA molecular species in WT (L) were quantified by ESI-MS/MS at 3 hpi. Data represent means ± SD (A) or ± SEM (C–L) from at least three independent experiments. Asterisks indicate statistically significant differences from mock by two-tailed Student’s *t*-test (\**P* < 0.05; \*\**P* < 0.01), except in (C), where one-way ANOVA with Fisher’s LSD post hoc test was used; different letters indicate statistically significant differences (*P* < 0.05).

Since PLA2α translocated to the apoplast upon *Pst AvrRpm1* inoculation (Figures 1B and 1C), it may generate lysophospholipids and free fatty acids from membrane phospholipids of either the host or invading pathogens, in the apoplast where it is activated by high Ca^2+^ concentration (Ryu et al., 2005). As expected, significant increases in lysophospholipids were detected 3 hpi with *Pst AvrRpm1* in Col-0 WT plant leaves compared to mock-inoculated plants (Figure 2C). Among the lipid products, LPE increased prominently, while lysophosphatidylglycerol (LPG) and LPC did not significantly increase. In contrast, *pla2α* mutant plants did not exhibit elevated LPE production in response to *Pst AvrRpm1* inoculation (Figure 2D). Notably, LPE18:1 was the most significantly increased LPE species in infected WT plants compared to mock-treated controls (Figure 2E). However, no increase in LPE as well as LPC and LPG was observed following inoculation with virulent *Pst* (Figure 2C), indicating that enhanced production of LPE is a specific response to avirulent *Pst*.

LPE18:1 was rarely detected in plant tissues (Figure 2E), suggesting a non-host origin. Given the strong antimicrobial activity of rPLA2α against *Pst* (Figure 1D) and the known ability of secretory PLA2 enzymes to hydrolyze microbial membranes, we next examined whether LPE18:1 may be derived from bacterial phospholipids. When lipid extracts of *Pst AvrRpm1* were incubated with rPLA2α protein, LPE but not LPC and LPG was predominantly generated (Figures 2F). Among the LPE species released from the bacteria, LPE18:1 was predominant, along with LPE16:0 (Figure 2G). To directly assess the origin of LPE species, *Pst AvrRpm1* was cultured with [¹⁴C]-ethanolamine to label bacterial PE and subsequently inoculated into leaves. At 0 h, radioactivity was predominantly in the PE fraction (with only ∼0.6% in LPE), whereas by 3 hpi ∼7% had shifted to the LPE fraction (Figure 2H), indicating that LPE is generated from bacterial phospholipids in planta.

We next asked whether LPE18:1 could also originate from host cell membrane phospholipids. Because PLA2α is secreted into the apoplast during ETI, it may hydrolyze plant cell membranes in addition to bacterial membranes. To test this possibility, rPLA2α protein was infiltrated into WT leaves, and LPE species released from plant cells were analyzed. Unlike *Pst AvrRpm1* inoculation, this treatment elevated levels of LPC and LPG, as well as LPE, at 3 hpi (infiltration) (Figures 2I and 2J). Moreover, LPE16:0 but not LPE18:1 was the major LPE species. These results suggest that LPE18:1, the major species of LPE increased during *Pst AvrRpm1* infection, is mainly released from the invading bacteria. In *Pst AvrRpm1*, PE constitutes ∼83% of total membrane phospholipids (Figure S2J). In *Pseudomonas*, the major fatty acids (FAs) incorporated into PE are 16:0, 16:1, and 18:1, whereas in *Arabidopsis*, PE species are predominantly composed of 16:0, 18:2, and 18:3 acyl chains, with 16:1 and 18:1 detected only at trace levels (Janse, 1991; Miquel and Browse, 1992). Free fatty acids (FFAs), another class of lipid products generated by PLA2α, also showed an overall increase with high variability following *Pst AvrRpm1* inoculation in WT plants, whereas no such increase was observed in the mutant plants (Figure 2K). The major FFAs detected in leaf tissues were stearic acid (18:0), palmitic acid (16:0), and linolenic acid (18:3), with only stearic acid exhibiting a significant increase upon *Pst AvrRpm1* infection (Figure 2L).

### Defects of the *pla2α* in ETI are restored by supplementation of LPE

Based on the phenotypes of *pla2α* plants compared with WT, we hypothesized that the immune defects of *pla2α* result from a failure to generate LPE18:1, which mediates downstream defense responses. To test this, we supplemented *pla2α* leaves with LPE18:1 after *Pst AvrRpm1* inoculation. LPE supplementation restored local immune responses in *pla2α*, suppressing avirulent bacterial growth to WT levels (Figure 3A). LPE specifically induced expression of defense-related genes *PR1* and *GST1*, but not wound-related genes *VSP1* and *JMT* (Figure 3B). Consequently, LPE treatment enhanced resistance to *Pst* infection, as evidenced by stronger defense responses in both WT and *pla2α* plants. LPE supplementation significantly reduced bacterial growth (Figure 3C) and alleviated disease symptoms (Figure 3D). Importantly, LPE itself had no direct antimicrobial effect on *Pst* growth in culture (Figure 3E), indicating that the observed resistance is not due to direct toxicity but rather to activation of host immune responses. LPE showed optimal activity at 2.4–4.0 nmol per leaf (Figure 3F), consistent with its function as a biologically active lipid signal in plant immunity. Together, these results demonstrate that LPE restores the immune defect of *pla2α* during *Pst AvrRpm1* infection and activates plant defense responses even in the absence of avirulent *Pst*. Consistent with this role in immune regulation, *pla2α* mutants also exhibited a modest but significant reduction in basal resistance to virulent *Pst*, although the defect was substantially more pronounced during ETI triggered by *Pst AvrRpm1* (Figure 3C).

**Figure 3.**
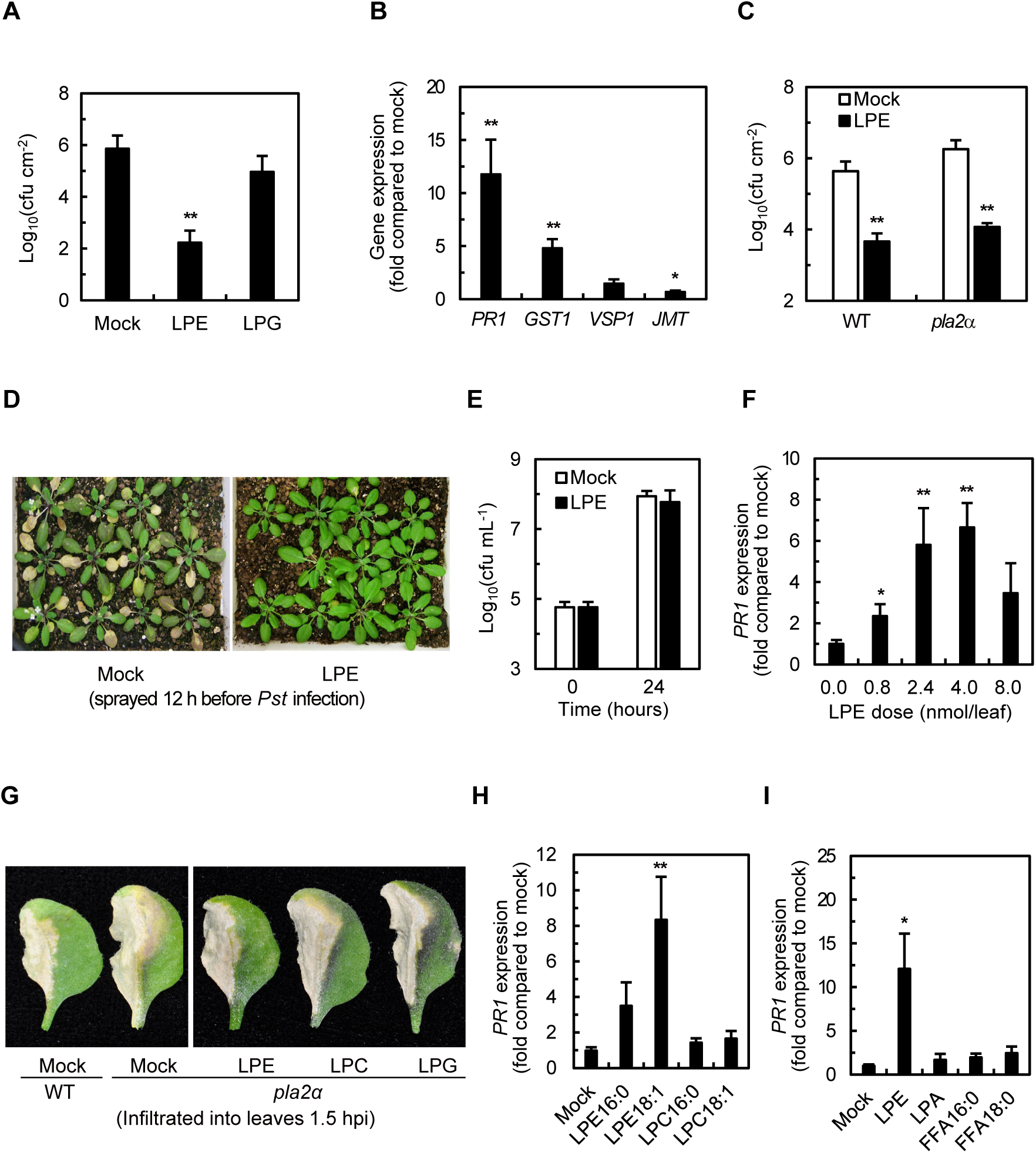
LPE18:1 activates local immune responses and restores ETI in *pla2α*. **(A)** Bacterial growth in *pla2α* plants pre-treated with lysophospholipids. Leaves were spray-inoculated with LPE18:1 or LPG16:0 (100 μM in 0.018% Silwet L-77) 6 h prior to spray inoculation with *Pst AvrRpm1* (4 × 10⁸ CFU mL⁻¹), and bacterial populations were quantified by CFU assay at 4 dpi. **(B)** Expression of defense- and wound-responsive marker genes in LPE18:1-treated WT plants. Plants were sprayed with LPE18:1 (100 μM in 0.018% Silwet L-77), and expression of *PR1*, *GST1*, *VSP1*, and *JMT* in leaves was quantified by RT-qPCR at 12 h post-treatment. Data are expressed as fold-change relative to mock-treated plants. **(C)** Bacterial growth in LPE18:1-pre-treated WT and *pla2α* plants. Leaves were sprayed with LPE18:1 (100 μM in 0.018% Silwet L-77) 6 h prior to spray inoculation with virulent *Pst* (1 × 10⁷ CFU mL⁻¹), and bacterial populations were quantified by CFU assay at 4 dpi. **(D)** Disease phenotypes of LPE18:1-pre-treated WT plants. Plants were sprayed with LPE18:1 (100 μM in 0.018% Silwet L-77) 12 h prior to spray inoculation with *Pst* (3 × 10⁸ CFU mL⁻¹) and photographed at 6 dpi. **(E)** Effect of LPE18:1 on *Pst* growth in culture. *Pst* cultures were supplemented with LPE18:1 or mock and incubated for 24 h, followed by CFU assay. **(F)** Dose-response relationship between LPE18:1 and *PR1* induction. Leaves were inoculated by applying four 10-μL droplets of LPE18:1 at the indicated doses per leaf. *PR1* expression was quantified by RT-qPCR at 12 h post-treatment. Data are expressed as fold-change relative to mock-treated plants. **(G)** MacroHR symptoms in *pla2α* plants supplemented with lysophospholipids. Leaves were syringe-infiltrated with *Pst AvrRpm1* (1 × 10⁷ CFU mL⁻¹), and the indicated lipids (100 μM in H_2_O) were syringe-infiltrated at 1.5 hpi. Plants were photographed at 6 dpi. **(H and I)** Specificity of *PR1* induction by LPE18:1 relative to other lipid classes and acyl species. Leaves were inoculated by applying four 10-μL droplets of the indicated lipids (100 μM each; equivalent to 4 nmol per leaf). *PR1* expression was quantified by RT-qPCR at 3 h post-treatment. Data are expressed as fold-change relative to mock-treated plants. Data represent means ± SD (A, C, and E) or ± SEM (B, F, H, and I) from at least three independent experiments. Asterisks indicate statistically significant differences from mock by two-tailed Student’s *t*-test (\**P* < 0.05; \*\**P* < 0.01).

These results support the hypothesis that the defect in *pla2α* is due to the failure to produce lipid mediators such as LPE, which trigger immune responses. Further supporting this, exogenous LPE18:1 application restored the ability of *pla2α* mutants to confine disease symptoms within the HR region. Notably, the boundary of HR lesion was visibly demarcated in the LPE-treated leaves, indicating effective spatial restriction of pathogen spread (Figure 3G). In contrast, application of LPC or LPG could not complement the phenotypic defect of the *pla2α* (Figure 3G). Neither LPC nor LPG was able to significantly induce *PR1* expression or suppress avirulent bacterial growth in the *pla2α* mutant to levels comparable to those induced by LPE (Figures 3A and 3H). LPC and LPG levels were not increased in response to avirulent *Pst* (Figure 2C), likely because PLA2α preferentially hydrolyzes PE over other phospholipids (Ryu et al., 2005), and because PC and PG are minor lipid components in *Pst* (Figure S2J).

These results underline the role of LPE as a primary signaling molecule that has unique functions in host immune responses. Supporting this notion, LPE, but not LPG or LPC, inhibits the activity of phospholipase Dα (PLDα) (Ryu et al., 1997), which is responsible for membrane deterioration. Furthermore, LPA, which has no headgroup, also failed to induce *PR1* expression (Figure 3I). Free fatty acids induced only modest *PR1* expression, much less than LPE at equivalent doses (Figure 3I).

### PLA2α and LPE mediate ICS1/NPR1-dependent salicylic acid signaling

ICS1 is localized in the plastid and plays a key role in host immune responses as a crucial enzyme in SA biosynthesis in *Arabidopsis* (Zhang et al., 2010; Shields et al., 2022; Zhu et al., 2024). The *ICS1* mutant *sid2* fails to express *PR1* in response to *Pst AvrRpt2* infection (Nawrath and Metraux, 1999; Liu et al., 2020; Yoo et al., 2022) or *Pst AvrRpm1* (Yoo et al., 2022). When *pla2α* plants were inoculated with *Pst AvrRpm1*, the expression of *ICS1* was significantly reduced compared to WT plants (Figure 4A). In contrast, *phenylalanine ammonia-lyase1* (*PAL1*) expression in infected *pla2α* plants was slightly increased. Consistent with this, the *pla2α* plants inoculated with *Pst AvrRpm1* produced lower levels of free SA than WT (Figure 4B). These results align with previous observations showing that the plastid-localized, ICS1-dependent isochorismate pathway is the main source of SA during ETI (Zhang et al., 2010; Shields et al., 2022). Thus, we hypothesized that in response to challenge of avirulent bacteria, *pla2α* is impaired in its ability to induce *ICS1* expression, resulting in reduced SA accumulation and compromised activation of SA-dependent downstream signaling required for *PR* gene expression. To examine this hypothesis, we asked whether exogenous LPE treatment induces *ICS1* expression. *Arabidopsis* leaves treated with LPE18:1 showed increased *ICS1* expression 6 h post-treatment (Figure 4C). In contrast, LPE treatment did not elevate *PAL1* expression. Induction of *ICS1* by LPE was followed by a two-fold increase in SA levels (Figure 4D). These results suggest that LPE elevates SA levels primarily through ICS1-mediated biosynthesis. The ICS1 dependency of LPE signaling was further confirmed by the observation that LPE failed to induce local immunity in *sid2* mutant plants (Figure 4E). LPE also failed to induce *PR1* expression in the *sid2* mutant (Figure 4F), as did *Pst AvrRpm1* inoculation (Yoo et al., 2022).

**Figure 4.**
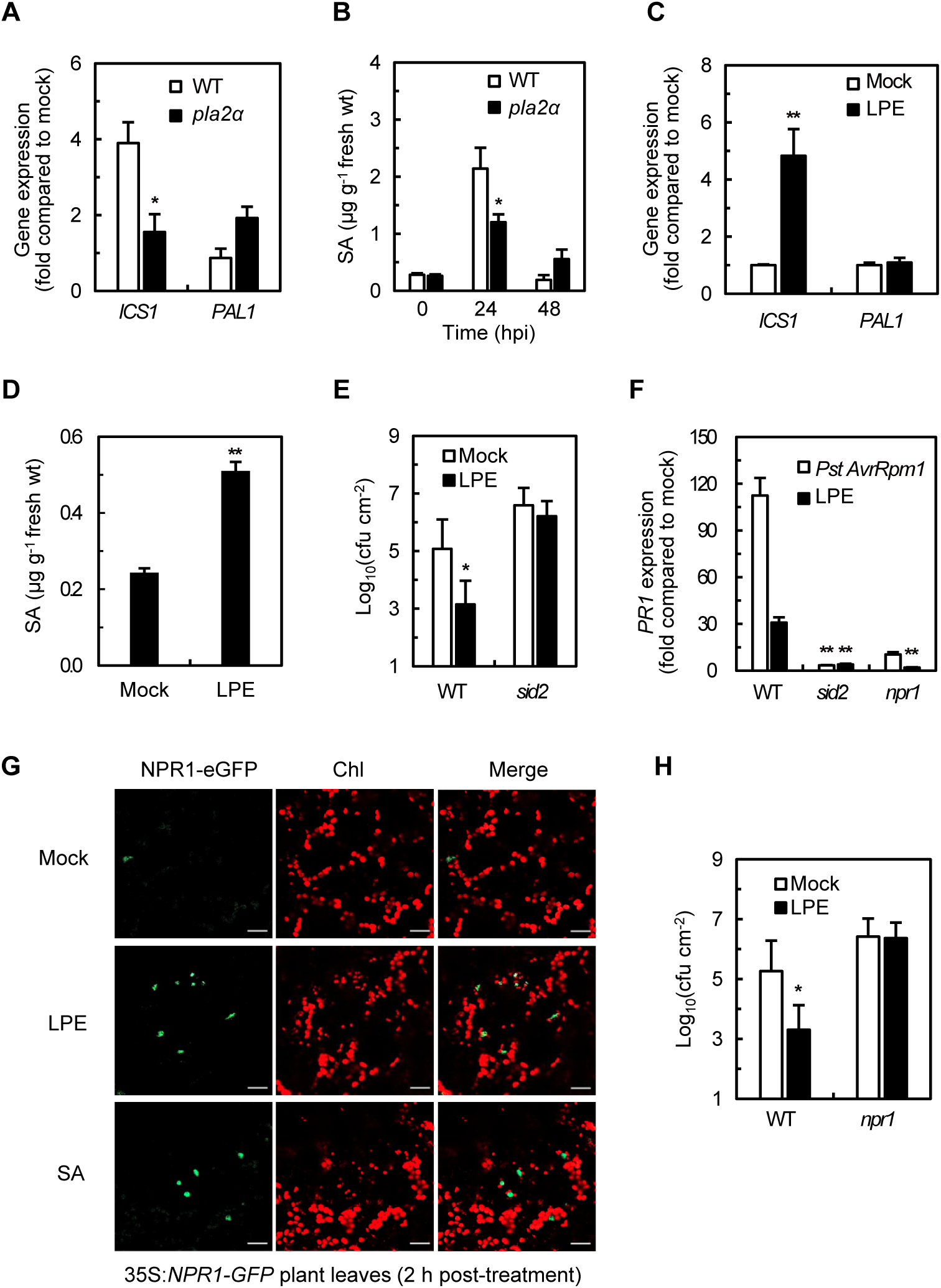
LPE18:1 activates local immunity through the ICS1–SA–NPR1 pathway. **(A)** Expression of *ICS1* and *PAL1* in WT and *pla2α* during ETI. Plants were spray-inoculated with *Pst AvrRpm1* (1 × 10⁸ CFU mL⁻¹), and gene expression in leaves was quantified by RT-qPCR at 8 hpi. Data are expressed as fold-change relative to mock-inoculated plants. **(B)** SA accumulation in WT and *pla2α* during ETI. Plants were spray-inoculated with *Pst AvrRpm1* (1 × 10⁸ CFU mL⁻¹), and SA levels in leaves were quantified by HPLC at the indicated hpi. **(C)** Expression of *ICS1* and *PAL1* in LPE18:1-treated WT plants. Leaves were inoculated by applying four 10-μL droplets of LPE18:1 (100 μM; equivalent to 4 nmol per leaf), and gene expression was quantified by RT-qPCR at 6 h post-treatment. Data are expressed as fold-change relative to mock-treated plants. **(D)** SA accumulation in LPE18:1-treated WT plants. Plants were spray-inoculated with LPE18:1, and SA levels in leaves were quantified by HPLC at 6 h post-treatment. **(E and H)** Bacterial growth in LPE18:1-pre-treated *sid2* and *npr1* plants. Plants were sprayed with LPE18:1 12 h prior to spray inoculation with virulent *Pst* (1 × 10⁷ CFU mL⁻¹), and bacterial populations were quantified by CFU assay at 4 dpi. **(F)** *PR1* expression in *sid2* and *npr1* following *Pst AvrRpm1* infection or LPE18:1 treatment. Plants were spray-inoculated with *Pst AvrRpm1*(1 × 10⁸ CFU mL⁻¹) or treated with LPE18:1 by applying four 10-μL droplets (100 μM; equivalent to 4 nmol per leaf). *PR1* expression was quantified by RT-qPCR at 12 h post-treatment. Data are expressed as fold-change relative to mock-treated plants. **(G)** LPE18:1-induced nuclear translocation of NPR1-GFP. Transgenic plants expressing *35S:NPR1-GFP* were treated with LPE18:1 (100 μM) or SA (500 μM) by spray inoculation and observed by confocal microscopy at 2 h post-treatment. Merged images show the overlap of NPR1-GFP fluorescence (green) and chlorophyll autofluorescence (red). Scale bar, 20 μm. Data represent means ± SEM (A–D, F, and H) or ± SD (E) from at least three independent experiments. Asterisks indicate statistically significant differences from mock by two-tailed Student’s *t*-test (\**P* < 0.05; \*\**P* < 0.01).

NONEXPRESSOR OF PR GENES 1 (NPR1) is a key regulator of SA-mediated immune responses, including *PR1* expression (Zhang et al., 2010; Liu et al., 2020). NPR1 activation requires its translocation from the cytoplasm to the nucleus due to SA-induced redox changes (Tada et al., 2008; Spoel and Dong, 2024; Zavaliev and Dong, 2024; Huang and Dong, 2025). Since NPR1 activation is mediated by SA, it is conceivable that LPE-induced SA accumulation activates NPR1. Application of LPE to transgenic 35S*:NPR1-GFP* plants induced nuclear translocation of NPR1, similar to SA treatment (Figure 4G). Moreover, LPE failed to induce *PR1* expression or confer resistance in the *npr1* mutant (Figures 4F and 4H). These data indicate that SA signaling is required for LPE-induced plant defense responses, with ICS1 and NPR1 functioning as key mediators. These results support the hypothesis that LPE promotes SA biosynthesis, leading to NPR1 activation and subsequent *PR1* expression.

### PLA2α-dependent transcriptomic changes in plant defense network

Transcriptional reprogramming is a core immune response in ETI that activates defense-related gene networks to enhance plant immunity (Bhandari and Brandizzi, 2024; Dodds et al., 2024). To define how PLA2α and its lipid product LPE shape defense-associated transcription during ETI, we performed RNA-seq analysis of *Arabidopsis* WT and *pla2α* plants treated with *Pst AvrRpm1*, LPE, or SA. Hereafter, up-regulated and down-regulated differentially expressed genes (DEGs) in each group are denoted by “+” and “−”, respectively.

More than half of the genes up-regulated in WT plants (WT**^+^**) in response to *Pst AvrRpm1* inoculation failed to be induced in the *pla2α* mutant (*pla2α*^+^). A similar ratio of genes that were down-regulated in WT (WT**^−^**) were not suppressed in the *pla2α* mutant (*pla2α***^−^**) (Figure 5A, Tables S1-S2). These results indicate that the *pla2α* mutation leads to a substantial reduction in the number of defense-related DEGs. To further identify genes most strongly affected in the *pla2α* mutant, which may play a core part of the *pla2α* deficit in disease responses, we compared gene expression between WT and *pla2α* plants after *Pst AvrRpm1* inoculation. This analysis revealed 676 up-regulated and 638 down-regulated genes in WT relative to *pla2α* (WT vs. *pla2α)* (Figures 5A and 5B), corresponding to 676 down-regulated and 638 up-regulated DEGs in *pla2α* relative to WT (*pla2α* vs. WT) (Figure 5C). As expected, 93% DEGs identified in the WT vs. *pla2α*⁺ comparison overlapped with WT⁺ DEGs (Figure 5A). Notably, 59% of these DEGs were also induced in *pla2α*, indicating that defense gene activation does occur in *pla2α* upon infection, albeit at substantially lower levels than in WT (Figure 5A).

**Figure 5.**
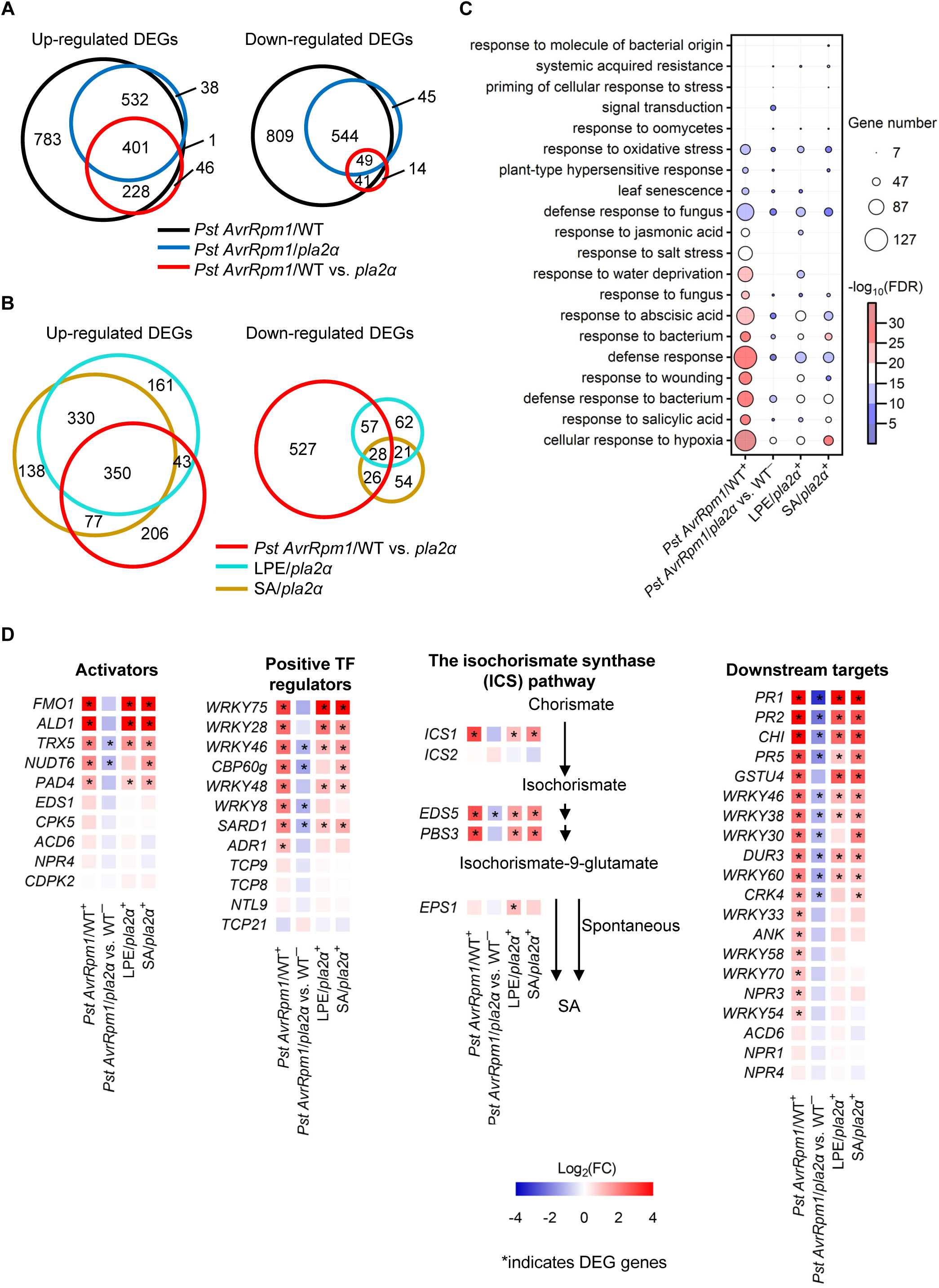
LPE18:1 restores the ETI transcriptional defense network impaired in *pla2α*. **(A)** Venn diagrams showing differentially expressed genes (DEGs) induced by *Pst AvrRpm1* infection in WT and *pla2α* plants relative to mock treatment, and DEGs identified by direct comparison of *Pst AvrRpm1*-infected WT versus *pla2α* plants. Plants were spray-inoculated with *Pst AvrRpm1* (1 × 10⁸ CFU mL⁻¹), and transcriptomes were profiled at 12 hpi by RNA-seq. **(B)** Venn diagrams showing DEGs in *pla2α* plants treated with LPE18:1 or SA, and their overlap with DEGs from the *Pst AvrRpm1*/WT versus *pla2α* comparison. **(C)** GO biological process enrichment analysis of DEG sets from (A) and (B). The top 15 enriched terms are shown. Bubble size represents the number of genes per term; color intensity represents statistical significance (−log_10_FDR). Upregulated DEGs (+) were analyzed for all comparisons except *Pst AvrRpm1*/*pla2α* vs. WT⁻, for which downregulated DEGs (−) were used. **(D)** Heatmaps of selected immune-related gene clusters across DEG sets in (C). Specific clusters include SA biosynthesis and signaling (*ICS1*/*2*, *EDS5*, *PBS3*, *EPS1*, *PR1*/*2/5*, and *CHI*), upstream ETI regulators (*FMO1*, *ALD1*, *EDS1*, and *PAD4*), and various WRKY transcription factors. Colors represent log_2_(fold change); asterisks indicate statistically significant DEGs (adjPval) ≤0.05). Data represent transcriptomic profiles from at least three independent biological replicates. See also Figures S3–S7 and Tables S1 and S2.

To assess whether the DEGs in WT vs. *pla2α* are regulated by LPE and SA, we compared DEGs among WT vs. *pla2α,* LPE-treated *pla2α* (LPE/*pla2α*), and SA-treated *pla2α* (SA/*pla2α*). Approximately 58% and 63% of DEGs in the WT vs. *pla2α*^+^ group were up-regulated by LPE and SA, respectively, indicating that both LPE and SA can restore a substantial portion of the down-regulated genes in the *pla2α* mutant, thereby contributing to the recovery of disease resistance in the *pla2α* mutant. Notably, over 76% of the DEGs up-regulated by SA and LPE were shared between the two treatments (Figure 5B), suggesting that LPE activates defense responses that substantially overlap with SA-responsive signaling. This finding supports the idea that LPE restores immunity in *pla2α* primarily by enhancing SA-mediated responses, with a smaller subset of genes (∼23%) reflecting SA-independent effects (Figures 5B and S3A). Furthermore, 350 DEGs were commonly up-regulated across all three treatments (*Pst AvrRpm1*, SA, and LPE), suggesting that they may represent a core regulatory module involved in host immune activation (Figure 5B). Analysis of WT plants treated with LPE and SA showed that approximately 36% and 38% of the WT+ genes were up-regulated by LPE and SA, respectively (Figure S3A). Most DEGs (>93%) up-regulated by LPE overlapped with the WT+ gene set, similar to the pattern observed with SA treatment (>99%) (Figure S3A).

To understand the functional categories of these DEGs, we performed Gene Ontology (GO), Kyoto Encyclopedia of Genes and Genomes (KEGG), and MapMan gene term analyses. GO analysis of biological process (BP) terms showed strong enrichment in defense-related categories such as “defense response to bacterium”, “defense response”, “response to fungus” and “plant-type HR” in the WT^+^ group, but these were dramatically reduced in the *pla2α*^+^ group (Figure 5C). These categories were significantly up-regulated following LPE or SA treatment. A similar pattern was observed in KEGG and MapMan analyses, where defense-related terms such as “Plant-pathogen interaction”, “Stress”, and “Stress.biotic”, were depleted in *pla2α* but restored by LPE or SA (Figures S4 and S5). In contrast, growth- and development-related terms down-regulated in WT were not significantly altered in *pla2α* and remained largely unresponsive to LPE or SA (Figures S6 and S7).

To further investigate whether PLA2α-induced immunity is predominantly dependent on SA-mediated signaling, we generated a heat map of well-characterized genes involved in SA biosynthesis, regulation, and downstream signaling (Gonzalez-Bayon et al., 2019; Huang et al., 2020). The heat map showed that genes in the ICS-dependent SA biosynthesis pathway, such as *ICS1*, *EDS5*, and *PBS3*, were suppressed in *pla2α* compared to WT in response to *Pst AvrRpm1* infection (Figure 5D). These genes were induced by LPE or SA. Similarly, the expression of transcription factors and activators involved in SA biosynthesis, including *FMO1*, *ALD1*, *TRX5*, *NUDT6*, *PAD4*, *WRKY8/28/46/48/75*, *CBP60g*, *SARD1*, and *ADR1*, was down-regulated in *pla2α* mutant but induced by LPE or SA (Figure 5D). Consequently, expression of downstream SA target genes, including defense markers such as *PR1/2/5*, *GSTU4*, *CHI*, *DUR3*, and *WRKY30/46*, was also reduced in *pla2α* but induced by LPE or SA. In conclusion, PLA2α-derived LPE induces transcriptomic changes important for ETI, primarily by activating SA-dependent defense signaling pathways.

### PLA2α-derived lipid signaling implicated in effector-triggered immunity

Together, our results consistently support the hypothesis that the impaired local immune responses observed in *pla2α* plants are attributable to the inability of secretory PLA2α to generate its lipid product LPE, which primarily activates SA-dependent defense signaling. Several additional lines of evidence further support this conclusion. RNA-seq analysis revealed that *PR1* expression did not differ significantly between WT and *pla2α* plants in response to LPE or SA treatment, in contrast to the marked difference observed following *Pst AvrRpm1* infection (Figure 6A). Exogenous application of rPLA2α to *pla2α* leaves induced the expression of *ICS1* and *PR1* (Figure 6B). In contrast, rPLA2α pre-inactivated by the PLA2 inhibitor manoalide failed to induce *PR1* expression (Figure 6C). In line with this, infiltration of manoalide into WT leaves prior to *Pst AvrRpm1* inoculation compromised the establishment of ETI (Figure 6D).

**Figure 6.**
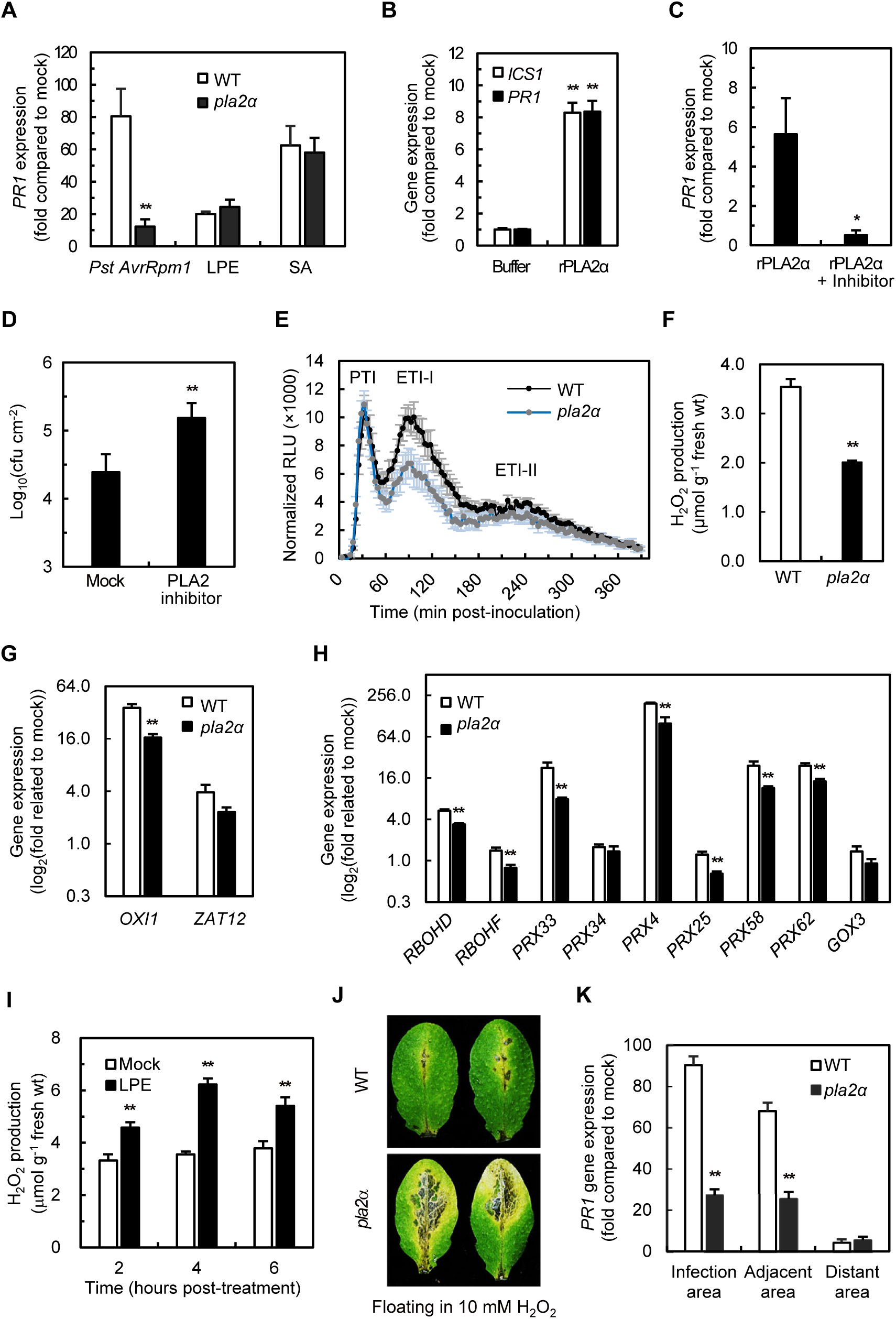
PLA2α–LPE signaling coordinates ROS homeostasis and spatially confined defense responses during ETI. **(A)** *PR1* expression in WT and *pla2α* under *Pst AvrRpm1* infection, LPE18:1, or SA treatment, derived from RNA-seq data (Figure 5). Data are expressed as fold-change relative to mock-treated plants. **(B and C)** Induction of *ICS1* and *PR1* by apoplastic PLA2α requires enzymatic activity. Leaves were syringe-infiltrated with active rPLA2α or manoalide-inactivated rPLA2α (rPLA2α + inhibitor), and gene expression was quantified by RT-qPCR at 6 h post-treatment. Data are expressed as fold-change relative to buffer-infiltrated plants. **(D)** Effect of PLA2 inhibition on bacterial growth during ETI. Leaves were syringe-infiltrated with manoalide prior to spray inoculation with *Pst AvrRpm1* (4 × 10⁸ CFU mL⁻¹), and bacterial populations were quantified by CFU assay at 4 dpi. **(E)** Time-course analysis of ROS production in WT and *pla2α* during PTI and ETI. Leaf discs were inoculated with *Pst AvrRpm1* (5 × 10⁷ CFU mL⁻¹) and ROS production was monitored for 390 min by luminol-based chemiluminescence assay. ETI-I and ETI-II denote early (60–165 min) and late (165–390 min) phases of ETI, respectively. RLU, relative luminescence units. **(F)** H_2_O_2_ accumulation in WT and *pla2α* during ETI. Leaves were syringe-infiltrated with *Pst AvrRpm1* (1 × 10⁷ CFU mL⁻¹), and H_2_O_2_ levels were quantified at 3 hpi by the ferric–xylenol orange (FOX) assay. **(G and H)** Expression of ROS-regulatory and ROS-producing genes in *pla2α* during ETI. Leaf discs were collected as in (E), and expression of ROS-regulatory genes (*OXI1* and *ZAT12*; G) and ROS-producing genes (*RBOHD*, *RBOHF*, *PRX33*, *PRX34*, *PRX4*, *PRX25*, *PRX58*, *PRX62*, and *GOX3*; H) was quantified by RT-qPCR at 1.5 hpi. The Y-axis is log_2_-transformed to facilitate visualization of large fold-change differences among genes. (**I**) H_2_O_2_ accumulation in LPE18:1-treated *pla2α* leaves. Leaves were infiltrated with LPE18:1 (100 μM) and H_2_O_2_ levels were quantified at the indicated time points by the FOX assay. (**J**) Sensitivity of WT and *pla2α* to exogenous H_2_O_2_. Leaves were floated on 10 mM H_2_O_2_ solution and photographed at 5 days post-treatment. (**K**) Spatial distribution of *PR1* expression relative to the *Pst AvrRpm1* inoculation site in WT and *pla2α*. Leaves were syringe-infiltrated with *Pst AvrRpm1* (1 × 10⁷ CFU mL⁻¹) and dissected at 8 hpi into three zones: infection area (infiltrated), adjacent area (2–3 mm from inoculation site), and distant area (>5 mm from inoculation site). *PR1* expression in each zone was quantified by RT-qPCR. Data are expressed as fold-change relative to mock-infiltrated plants. Data represent means ± SEM (A–C, E–H, K) or ± SD (D, I) from at least three independent experiments. Asterisks indicate statistically significant differences from mock or WT by two-tailed Student’s *t*-test (\**P* < 0.05; \*\**P* < 0.01), except in (C), where a one-tailed Student’s *t*-test was used.

Given the partial defect in HR execution observed in *pla2α* leaves and the close association between HR and ROS, we examined the relationship between PLA2α and ROS during *Pst AvrRpm1*-triggered immunity. Luminol-based assays using leaf discs revealed that WT and *pla2α* plants exhibited comparable ROS bursts during pattern-triggered immunity (PTI), whereas ETI-associated ROS peaks (Yuan et al., 2021; Hong et al., 2023), particularly the early ETI-I phase, were markedly reduced in *pla2α*, indicating compromised ROS production during ETI (Figures 6E, 6F, and S8A). Consistent with this defect, the expression of both ROS-regulatory and ROS-producing genes was significantly reduced in *pla2α* (Figures 6G, 6H, and S8B), a trend further supported by RNA-seq analysis (Figures S8C and S8D). Importantly, treatment with either LPE18:1 or SA induced the expression of these ROS-related genes (Figures S8E and S8F). Additionally, LPE18:1 treatment promoted ROS production (Figures 6I), consistent with previous reports that SA can stimulate ROS generation (Chen et al., 1993; Torres et al., 2002; Mammarella et al., 2015), although complex feedback interactions between SA and ROS have also been reported (Torres et al., 2002). These findings indicate that PLA2α–LPE signaling, acting through the SA pathway, is required for proper ROS homeostasis during HR execution. Consistently, whereas *sid2* mutants exhibited HR defects similar to those of *pla2α*, *npr1* mutants retained a WT-like HR phenotype following *Pst AvrRpm1* inoculation (Figure S2G), suggesting that ICS1-dependent SA biosynthesis contributes to HR development at infection sites, whereas NPR1-mediated signaling is less critical for this process.

In neighboring cells adjacent to the infection site, ROS containment is critical for preventing further expansion of HR (Ross, 1961; Costet et al., 1999; Jacob et al., 2023a; Jacob et al., 2023b). The failure to spatially restrict HR in *pla2α* mutants (Figures 1E and 3G) indicates impaired HR containment, as further demonstrated by floating-leaf assays. *pla2α* leaves exhibited increased cell damage compared to WT in response to exogenous H_2_O_2_ (Figure 6J), consistent with a reduced detoxification capacity. This phenotype coincides with impaired induction of genes involved in ROS regulation and detoxification, including *OXI1*, *ZAT12*, *CAT2*, and *APX1*, in the mutant (Figures 6G, S8B, and S8C). Importantly, LPE18:1 treatment induced the expression of *OXI1* and *ZAT12* (Figure S8F) and rescued the associated phenotypic defects (Figure 3G).

To further assess spatial defects in defense gene activation, we analyzed *PR1* expression in three tissue zones: the *Pst AvrRpm1*-infected area, an adjacent area, and a distant area. *PR1* expression was significantly reduced in both infected and adjacent regions of *pla2α* plants compared to WT, but not in distant tissues at 8 hpi (Figure 6K). Notably, while low levels of bacterial colonization were detected in non-infected regions of WT leaves, colonization in these regions was significantly elevated (∼2.5-fold) in *pla2α* mutants (Figure S8G), indicating that PLA2α–LPE signaling limits bacterial establishment and proliferation in tissues adjacent to infection sites. Collectively, these results demonstrate that PLA2α–LPE signaling is required not only for HR execution at infection sites but also for HR containment, spatial reinforcement of defense gene expression, and restriction of bacterial spread and subsequent colonization in neighboring cells.

Based on these findings, we propose a working model for PLA2α-mediated lipid signaling during ETI (Figure 7). Upon inoculation with avirulent *Pst*, *PLA2α* is transiently and moderately induced in an effector-triggered manner. Secreted PLA2α accumulates in the apoplast, where it hydrolyzes bacterial phospholipids to release LPE, predominantly LPE18:1, from invading pathogens. This apoplastic lipid signal promotes ICS1-dependent SA accumulation and reinforces downstream immune outputs, including ROS homeostasis and transcriptional reprogramming. In infected cells, these processes enable proper execution of the HR; in neighboring cells, LPE acting as a short-range signal reinforces defense gene expression and helps confine the HR boundary. Together, PLA2α-derived LPE18:1 coordinates SA signaling, balanced ROS dynamics, spatial control of HR, and robust defense gene expression, ultimately reinforcing local immunity during ETI (Figure 7).

**Figure 7.**
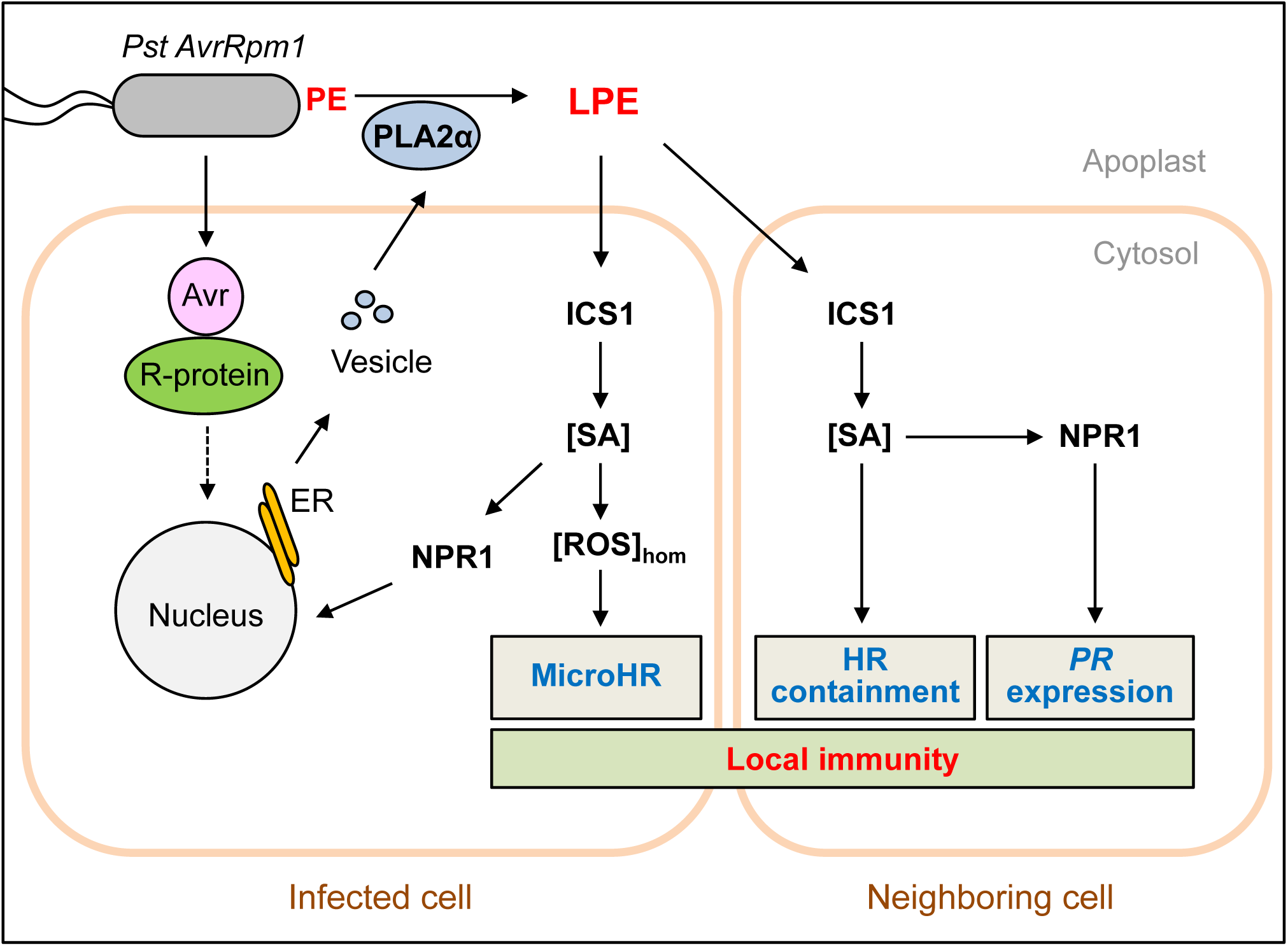
Proposed model for PLA2α–LPE18:1-mediated local immune signaling during ETI. Recognition of the bacterial effector AvrRpm1 by the NLR immune receptor RPM1 triggers transient induction of secretory PLA2α. Newly synthesized PLA2α is secreted into the apoplast, where it hydrolyzes phosphatidylethanolamine (PE) derived predominantly from invading bacterial membranes to generate LPE18:1. LPE18:1 acts as a short-range lipid immune signal that (i) activates ICS1-dependent SA biosynthesis, (ii) drives NPR1-dependent transcriptional reprogramming including *PR* gene expression, (iii) promotes ROS homeostasis, and (iv) ensures spatially contained HR at the infection focus (microHR). Together, these outputs coordinate local immunity and restrict pathogen spread to surrounding cells.

## DISCUSSION

A central unresolved question in plant immunity concerns the identity of molecular signals that bridge intracellular NLR-mediated effector recognition with downstream ETI execution. Our study addresses this gap by identifying pathogen-derived LPE18:1 as a previously undescribed class of ETI-mediating lipid signal. Canonical pattern-triggered immunity (PTI) signals, such as flg22 and chitin fragments, are derived from pre-existing pathogen structural components (Jiang et al., 2026). Chitin fragments, in particular, are enzymatically released from fungal cell walls by constitutively expressed host chitinases and perceived by cell-surface pattern recognition receptors (PRRs). In this regard, LPE18:1 shares a conceptual parallel with chitin-derived elicitors in that both are generated from microbial structural components through host enzymatic activity.

A key distinction, however, lies in the upstream activation requirement and the biochemical nature of signal generation. PTI can also involve host enzyme–mediated processing of pathogen-derived substrates—as exemplified by the apoplastic subtilases SBT5.2 and SBT1.7, which cleave bacterial flagellin to release the immunogenic flg22 epitope (Matsui et al., 2024), and by SBT3.3, a PTI-induced subtilase that amplifies immune responses through SA-dependent chromatin remodeling rather than direct substrate processing (Ramirez et al., 2013). However, these activities act on already-recognized PAMPs and do not require prior NLR activation. In contrast, LPE18:1 is generated from immunologically inert bacterial membrane phospholipids through ETI-induced, PLA2α-mediated hydrolysis—a process strictly gated by intracellular NLR recognition. LPE18:1 thus represents a mechanistically distinct category: an ETI-gated lipid mediator arising from host enzymatic remodeling of pathogen structural components during effector-triggered immune activation.

We further demonstrate that secretory PLA2α and its lipid product, LPE, reinforce NLR-triggered local immune responses primarily through the activation of *ICS1*-dependent SA signaling and coordinated regulation of ROS homeostasis. The apoplastic localization of LPE is well suited for rapid short-range signal propagation, consistent with the known ability of lysophospholipids to traverse plant membranes within minutes (Viehweger et al., 2002). We propose that PLA2α-generated LPE serves as a localized signal that potentiates immunity at the site of pathogen ingress and thereby spatially restricts HR–associated cell death. Moreover, the use of pathogen-derived LPE18:1 provides plants with a chemically distinct molecular proxy of infection. Because *Pseudomonas* and many Gram-negative bacterial membranes are enriched in 18:1-containing phosphatidylethanolamine species relative to plant membranes (Janse, 1991; Miquel and Browse, 1992), LPE18:1 functions as a pathogen-associated "lipid signature" that enables site-restricted immune activation precisely at infection sites.

Our findings support the hypothesis that PLA2α secretion into the apoplast is essential for initiating PLA2α-mediated lipid signaling in immune responses. In a complementary experiment, expression of secretory PLA2α in the *pla2α* mutant restored the immune defects, but expression of a non-secreted PLA2α (lacking the signal peptide) in the *pla2α* mutant failed to rescue the immune defects (Figures 1J and 1K), emphasizing the importance of apoplastic localization for PLA2α function. Consistent with this hypothesis, exogenous supply of rPLA2α, but not manoalide-inactivated rPLA2α, to the apoplast of *pla2α* leaf tissues induced defense gene expression (Figures 6B and 6C) by generating LPE16:0 instead of LPE18:1 (Figure 2J). Additionally, treatment of WT leaf tissues with the PLA2 inhibitor manoalide prior to bacterial infection suppressed the defense response to *Pst AvrRpm1* (Figure 6D), indicating that active PLA2 enzymatic activity in the apoplast is necessary for triggering host immunity. Furthermore, immune defects in *pla2α* were more evident in mature leaves, where PLA2α localizes to the apoplast, than in juvenile leaves, where it remains in the ER/Golgi as shown in the original image of Figure S2H. These results are consistent with a previous report (Froidure et al., 2010) that PLA2α only slightly or negatively regulates *Arabidopsis* resistance under short-day conditions, which extend the juvenile phase. Under short-day conditions we also observed that the immune phenotype of *pla2α* was substantially attenuated, consistent with the observations reported by Froidure et al. (2010). These findings highlight the apoplast as a critical battleground between plant hosts and invading pathogens (Wan et al., 2021; Del Corpo et al., 2024).

An important feature of ETI is the robust production of ROS, which function as both antimicrobials and signaling mediators. Later in ETI, *pla2α* mutants exhibited defective HR, which was underpinned by impaired ROS-related transcriptional reprogramming. This included reduced expression of *RBOHD*, a plasma membrane NADPH oxidase responsible for apoplastic ROS generation; *class III peroxidases* (e.g., *PRX33/34*), which modulate ROS levels and mediate cell wall strengthening; and *APX1*, a key ROS-scavenging enzyme that maintains redox homeostasis. Most of the defects were rescued by LPE or SA treatment, suggesting that PLA2α–LPE signaling coordinates both ROS-generating and scavenging modules to stabilize redox homeostasis and enable proper HR execution (Mammarella et al., 2015; Waszczak et al., 2018; Kaur et al., 2021; Ngou et al., 2021; Yuan et al., 2021; Ugalde, 2023; Cao et al., 2024). While our results suggest that LPE contributes to maintaining redox balance, further detailed analyses of ROS dynamics will be valuable to clarify how PLA2α–LPE signaling maintains ROS homeostasis during ETI.

Multiple lines of evidence confirm that LPE treatment restores immunity in *pla2α* mutants through specific biological effects, not nonspecific detergent action. Among the lysophospholipids tested, only LPE, not LPA, LPC or LPG, complemented the immune defects in *pla2α*, suggesting a specific and selective role of LPE in plant defense. More than 93% of LPE-induced genes overlapped with those induced by *Pst AvrRpm1*, and these were predominantly defense-related. Notably, LPE selectively induced defense-related genes but not general stress-responsive ones. LPE-induced SA signaling depended on ICS1 and NPR1, as does NLR-mediated SA signaling during infection by avirulent bacteria. The induction of *PR* genes was triggered by a small amount of LPE (2–4 nmol per leaf), which is only 10–20 times higher than the *in vivo* LPE increase upon *Pst AvrRpm1* inoculation (Figure 3F). This requirement for higher exogenous LPE concentrations is consistent with partial loss of bioactive molecules during penetration through the waxy cuticle and leaf tissues. Notably, the concentration of 100 μM, which is effective for plant treatment, is comparable to physiological levels of LPE in human plasma (10–50 μM) (Tan et al., 2020). Like other signals, LPE showed optimal bioactivity at 4 nmol per leaf (40 μL of 100 μM), with effects declining at higher doses (>8 nmol per leaf). Our results are consistent with apoplastic LPE being low at baseline but locally enriched in LPE18:1 at infection foci, a pattern that could underlie strong signaling despite modest whole-leaf changes.

Collectively, our study strongly indicates that LPE fulfills the criteria for a local immune signal in plants for several compelling reasons. First, the rapid ETI-induced secretion of PLA2α leads to immediate LPE generation in the apoplast. Second, LPE—whether endogenous or exogenous—provokes robust transcriptional reprogramming that confers resistance against both virulent and avirulent *Pst*. Third, LPE alone is sufficient to elicit immune responses in the absence of pathogen effectors. Fourth, downstream signaling of PLA2α–LPE mirrors the well-characterized ICS1–SA-NPR1 pathway. Finally, the chemical uniqueness of bacterial LPE18:1, which is nearly absent in host cells but introduced upon infection, optimizes its function as a distinct immune elicitor. These findings align with emerging models in which small molecules and lipid-derived signals coordinate complex immune networks (Boller and Felix, 2009; Jiang et al., 2026). Consistent with the concept of localized acquired resistance (Jacob et al., 2023a; Nobori et al., 2025), we propose that LPE generated at infection sites diffuses into neighboring cells to reinforce spatially confined defense responses in coordination with H_2_O_2_ signaling.

The identification of pathogen-derived LPE18:1 as an ETI-mediating immune signal raises the possibility of its application as a potent plant immune inducer. While numerous immune-activating agents—including PTI-associated elicitors such as chitin, chitosan, and flg22—have been explored as plant defense activators, these typically confer only a moderate level of basal resistance. In contrast, LPE18:1 is generated specifically through ETI-induced PLA2α activity and is capable of reconstituting key immune outputs of ETI—including ICS1-dependent SA biosynthesis, NPR1-mediated transcriptional reprogramming, and ROS homeostasis—elevating resistance to levels comparable to those achieved during avirulent *Pst* infection. This capacity to reconstitute ETI-level resistance through a single lipid-derived signal, effectively bypassing the requirement for upstream NLR-mediated effector recognition, distinguishes LPE18:1 from canonical PTI elicitors and synthetic activators such as benzothiadiazole (BTH), which engage immune pathways only partially or indirectly. As a sustainable and potent bio-based activator—a conceptual "plant vaccine"—LPE18:1 provides a biologically grounded strategy to achieve robust and spatially confined disease resistance.

We propose the following model of ETI-associated local immunity under natural infection conditions. Avirulent *Pst* enters *Arabidopsis* leaves primarily through stomata and frequently elicits microHR lesions near stomata and trichomes. During early ETI, PLA2α is secreted to the apoplast and generates LPE18:1 from invading bacterial phospholipids. As a pathogen-derived lipid signal, LPE18:1 promotes transcriptional reprogramming and ROS homeostasis required for proper microHR execution in infected cells, and reinforces defense activation in neighboring cells—thereby driving HR containment and *PR* gene induction. This tight spatial control confines defense responses to the infection site, minimizes collateral damage to surrounding healthy tissues, and ultimately establishes local immunity. Consequently, infected leaves largely recover, leaving only small, localized microHR lesions that resolve over time. In the *pla2α* mutant, however, the lack of LPE18:1 production compromises both microHR execution and HR containment. This dual defect manifests as failure to establish distinct HR boundaries, allowing lesions to expand beyond initially infected sites and spread into surrounding tissues. Under these conditions, avirulent *Pst* is no longer efficiently restricted and can proliferate extensively, resulting in disease symptoms that resemble virulent infection, particularly evident in spray-inoculated older leaves. Importantly, proper microHR formation requires ICS1 but is largely independent of NPR1. Nevertheless, uncontrolled bacterial proliferation is severe in both the *sid2* and *npr1* mutants (Cao et al., 1994; Wildermuth et al., 2001), underscoring the essential contribution of the ICS1–NPR1–PR signaling circuit to effective local immunity. These results demonstrate that PLA2α-mediated LPE18:1 production is essential for spatially controlled ETI responses, orchestrating both HR execution at infection sites and HR containment in surrounding tissues through coordinated regulation of ROS dynamics and ICS1–NPR1–PR signaling (Figure 7). Despite this advance, the molecular mechanisms underlying LPE18:1 perception and the upstream events linking NLR activation to PLA2α secretion remain important open questions. Addressing these gaps will deepen our understanding of how lipid-derived signals are integrated into plant immune networks to orchestrate spatially confined defense responses.

Our findings are also relevant to animal systems, where lipid signaling in immunity is a growing area of research (Tan et al., 2020; Peng et al., 2021; Murakami et al., 2023). Similar to findings in animal PLA2α, *Arabidopsis* PLA2α plays dual roles: both as a scavenger by killing invading pathogens and as a generator of signal molecules such as LPE that activate downstream defense-related signaling. In contrast, in animals, lysophospholipids including LPA, LPI, and LPS are established signaling mediators, whereas LPE has not yet been implicated in this role (Tan et al., 2020; Kano et al., 2022; Chakraborty and Kamat, 2024). By demonstrating LPE’s signaling function in plants, our work introduces a concept that could open new avenues for mammalian systems as well. *Arabidopsis* secretory PLA2α has low sequence homology to animal secretory PLA2s (sPLA2s), yet key domains such as the Ca²⁺-binding loop and catalytic motifs are conserved (Figure S9 and Table S3). Like *Arabidopsis* PLA2α (Ryu et al., 2005), human sPLA2-IIA also prefers PE as a substrate (Murakami et al., 2023). Since PE accounts for ∼83% of bacterial membrane phospholipids, this substrate preference may reflect a conserved evolutionary strategy for pathogen recognition. Similar to our findings in *pla2α*, the immune deficiency in sPLA2-V-deficient human macrophages was restored by LPE treatment following IL-4 stimulation, which normally enhances phagocytosis and sPLA2-V expression (Rubio et al., 2015). Nevertheless, our understanding of how sPLA2-derived lysophospholipids, such as LPE, function in mammals remains limited, partly due to the incomplete identification of their enzymatic sources. Our findings in *Arabidopsis* may thus offer a useful framework for probing analogous lipid-signaling mechanisms in mammalian immunity.

In conclusion, our findings establish a previously unrecognized lipid-mediated immune signaling cascade in which secreted PLA2α and its product LPE act as key regulators of ETI. By establishing LPE as an apoplastic signal capable of coordinating responses at infected sites, we provide new mechanistic insight into how lipid-derived cues orchestrate local immune responses. Given the evolutionary conservation of secretory PLA2 enzymes and lysophospholipid signaling across eukaryotes, these insights not only advance our understanding of plant immunity but also raise the possibility that pathogen-derived lipid signaling represents a broadly conserved immune strategy—one that may extend to human innate immunity and warrants future investigation across kingdoms.

## RESOURCE AVAILABILITY

### Lead contact

Further information and requests for resources and reagents should be directed to the lead contact, Stephen B. Ryu.

### Materials availability

Plasmids and plant lines generated in this study are available from the lead contact upon request.

### Data and code availability

The RNA-seq raw data reported in this paper have been deposited in the Korea Sequence Read Archive (KRA) at the Korea Bioinformation Center (accession number: KAP241514). This paper does not report original code.

## ACKNOWLEDGEMENTS

We express our gratitude to Hae Jin Kim for her contribution. This work was supported by grants from the Plant Diversity Research Center of the 21st Century Frontier Research Program (PF0330502; PF06305-01) funded by Korea MEST; from the 2nd BioGreen 21 Program (PJ009029; PJ011091; PJ013486) funded by Korea RDA; from the NRF (RS-2024-00445024; RS-2025-00519975) funded by Korea MSIT, and in part by the KRIBB Project Initiative Program (KGM1082511).

## AUTHOR CONTRIBUTIONS

S.B.R. conceived and designed the research with input from H.Y.L.; H.Y.L. performed the initial experiments with assistance from S.J. for SA quantification. J.J., K.K., and T.L. independently replicated and validated key experiments to ensure reproducibility. T.L. conducted transcriptomic and bioinformatic analyses, and S.K. performed supplementary experiments. S.B.R. carried out lipidomic profiling and radioisotope studies. K.B. and I.H. provided critical intellectual input throughout the study. S.B.R. wrote the manuscript with contributions from T.L. and S.K. All authors reviewed and approved the final manuscript.

## DECLARATION OF INTERESTS

S.B.R. is an inventor on patent applications filed by the Korea Research Institute of Bioscience and Biotechnology related to this work, covering technologies for enhancing both plant and human innate immunity through phospholipase-mediated lipid signaling. T.L. and S-C.K. are employed by Ourbio Co., Ltd. and JICO Ltd., companies that have a potential commercial interest in the findings described in this manuscript. The remaining authors declare no competing interests.

## DECLARATION OF GENERATIVE AI AND AI-ASSISTED TECHNOLOGIES IN THE WRITING PROCESS

During the preparation of this work, the authors used Claude and ChatGPT in order to ensure that there are no grammatical errors in the manuscript. After using this tool/service, the authors reviewed and edited the content as needed and take full responsibility for the content of the publication.

## EXPERIMENTAL MODEL AND STUDY PARTICIPANT DETAILS

### Plant materials and growth conditions

The plants were grown at 22°C, 60% relative humidity, with a 16 h photoperiod, and a photon flux density of 120 μmol m^-2^ s^-1^. The fully mature leaves from 4-week-old plants were used for most assays, as defense responses are age-dependent. *A. thaliana* mutants including *pla2α* (Salk_099415), *pla2α*-*II* (CS857021), *sid2* (Salk_042603), *and npr1-5* (CS3724) were obtained from the Arabidopsis Biological Resource Center (ABRC). The generation of transgenic *Arabidopsis* lines co-expressing both *PLA2α-DsRed2* and *ST-GFP* was performed as previously described (Lee et al., 2010). The *NPR1-GFP* transgenic line was provided by X. Dong (Duke University, USA). To avoid confusion with animal isoforms, we adopt the flat-text format (PLA2α) for secretory phospholipase A2α, which has previously been designated as PLA_2_α in plant studies.

### Complementation and transgenic line construction

Mutant *pla2α* plants were complemented by transformation with the floral dip method using a binary plant transformation vector, the pCAMBIA1300 clone, containing *PLA2α* genomic DNA bearing the native promoter region (-1175 bp to +922 bp), inserted at the *BamHI* site.

For histochemical localization, a *PLA2α*-promoter*:GUS* (pPLA2α*:GUS*) construct was created by cloning the *PLA2α* promoter region (-1175 bp to +3 bp relative to the ATG start codon) into the *HindIII* and *BamHI* sites of the pBI101 vector (Jung et al., 2012). Primer sequences used for cloning are listed in Table S4. To redirect the subcellular localization of PLA2α from the apoplast to the cytosol, a modified version of the pPLA2α*:PLA2α* construct (*PLA2ΔSP*) was constructed by deleting the signal peptide sequence from the complementary vector mentioned above.

### Bacterial inoculation of plants

*Pseudomonas syringae* pv. *tomato* DC3000 (*Pst*) and its avirulent strains expressing *AvrRpm1*, *AvrRpt2*, and *AvrRps4* were used in this study. The *Pst AvrRpm1* strain was provided by Y.J. Kim (Korea University, Seoul), *Pst AvrRpt2* by J.M. Park (KRIBB, Daejeon), and *Pst AvrRps4* by R. Innes (Indiana University, USA). Bacterial strain culture and handling followed the protocol described (Katagiri et al., 2002). King’s B medium was used for *Pst* strains culture supplemented with appropriate antibiotics as follows: rifampicin (50 μg mL^-1^) for *Pst* DC3000, rifampicin (50 μg mL^-1^) and kanamycin (50 μg mL^-1^) for *Pst AvrRpm1*, *Pst AvrRpt2*, and *Pst AvrRps4*. Unless otherwise specified, bacterial inoculation was performed by spray to mimic natural infection through stomatal entry. Syringe-infiltration was used in selected experiments requiring synchronized and spatially confined infection.

For gene expression analysis with qRT-PCR, *Arabidopsis* plant leaves were inoculated either spraying with a bacterial suspension (1 × 10^8^ CFU mL^-1^ in 0.018% (v/v) Silwet L-77) or by syringe-infiltration with a bacterial suspension (1 × 10^7^ CFU mL^-1^ in H_2_O).

Bacterial growth assays were conducted by spray inoculation of plants with *Pst* or *Pst AvrRpm1* at 1 × 10^7^ or 4 × 10⁸ CFU mL^-1^, respectively, in 0.018% (v/v) Silwet L-77 and 7 mM MgCl_2_. After spray-inoculation, plants were air-dried and then loosely covered with transparent plastic to maintain high humidity without visible dew formation, which enhanced bacterial proliferation and the differences between genotypes. At the indicated time points, 6 leaf discs (5 mm diameter) were collected per sample from three independent plants and homogenized using a TissueLyser II (Qiagen, Germany). Homogenates were serially diluted in sterile distilled water, and 10-μL aliquots of each dilution were plated onto KB agar medium supplemented with appropriate antibiotics. Colonies were counted after incubation at 28°C for 48 h and expressed as log₁₀(CFU cm⁻²).

### Protein subcellular localization

Subcellular localization of ST-GFP and PLA2α-DsRed2 in *Arabidopsis* leaves was examined using a confocal laser scanning microscope (LSM780, Zeiss). Cell walls were stained with 0.1 mg mL^-1^ Fluorescent Brightener 28 (FB-28, Sigma-Aldrich). Fluorescence signals were captured using the following excitation/emission settings: GFP, 488/505–530 nm; DsRed2, 543/560–615 nm; FB-28, 405/425–475 nm (Jung et al., 2012; Yin et al., 2021). Leaves from 35S*:NPR1-GFP* transgenic plants were imaged 2 h after treatment using the same settings. Chlorophyll (Chl) autofluorescence was detected with excitation at 488 nm and emission at 645–700 nm.

### Protein fractionation and extraction, MDH activity assay and Western blotting

Protein fractionation was performed as previously described (Nakano et al., 2020). Total, apoplastic, and intracellular protein fractions were isolated from the leaves of transgenic *Arabidopsis* plants co-expressing ST-GFP (Golgi marker) and PLA2α-DsRed2 (Lee et al., 2010) at 3–4 h post-inoculation (hpi) following spray treatment with either mock solution (dH_2_O with 0.018% Silwet L-77) or *Pst AvrRpm1* (1 × 10⁸ CFU mL^-1^ in mock solution).

Twelve to fifteen whole plants per treatment were vacuum-infiltrated with ice-cold infiltration buffer (5 mM sodium acetate, 0.2 M calcium chloride, pH 4.3). After gently blotting the plants with paper towels, they were inserted into a 20 mL blunt-end needleless syringe placed within a 50 mL conical tube and centrifuged at 1,000 ×g for 20 min at 4°C. The bottom fraction was collected as the apoplastic protein fraction (AF). Total protein fractions (TF) and intracellular protein fractions (ICF) were extracted from leaves before and after centrifugation, respectively, using a protein extraction buffer containing 50 mM Tris-HCl (pH 7.5), 10% glycerol, 5 mM EDTA, 5 mM DTT, 150 mM NaCl, and 1x protease inhibitors (cOmplete™, Roche).

To assess cytoplasmic contamination in the AF, MDH activity was measured in AF and TF fractions using a modified protocol (Husted and Schjoerring, 1995). A 25 µL aliquot of each fraction was added to 1 mL of assay buffer containing 100 mM Tris-HCl (pH 7.4), 0.094 mM β-NADH-Na_2_, and 0.17 mM oxaloacetic acid. MDH activity was monitored spectrophotometrically by the decrease in A_340_ (OPTIZEN Alpha, K Lab Co., Ltd.). PLA2α-DsRed2 was detected by Western blot using rabbit anti-DsRed2 antibody (1:1,000; Takara) and goat anti-rabbit IgG-HRP secondary antibody (1:5,000; Santa Cruz Biotechnology). Signals were visualized with Clarity™ Western ECL Substrate (Bio-Rad) and imaged using the ChemiDoc Imaging System (Bio-Rad). Band intensities were quantified using Image Lab 6.1 (Bio-Rad).

### Lipid extraction from leaves and ESI-MS/MS analysis

Total lipids were extracted as previously described (Ryu and Wang, 1996) from *Arabidopsis* leaves (6 leaves from 3 plants for each sample) sprayed with mock or *Pst AvrRpm1* suspension (1 × 10^8^ CFU mL^-1^ in 0.018% Silwet L-77).

Individual phospholipids and FFAs were extracted following the protocol (Ryu and Wang, 1998). *Arabidopsis* leaves were incubated in preheated isopropanol (75°C, containing 0.01% BHT) for 15 min. Subsequently, chloroform and water were added (isopropanol:chloroform:water = 3:1:0.5, v/v/v) at room temperature, and the mixture was shaken continuously for 1 h. After transferring the supernatant, the tissue was extracted three additional times with a solvent mixture of chloroform:methanol (2:1, v/v, containing 0.002% BHT), shaking for 30 min each time. Combined extracts (approximately 10–12 mL) were washed by adding 1.0 mL of 2 M KCl, mixed thoroughly, and centrifuged at 3,000 rpm for 5 min. The upper aqueous phase was removed, followed by a wash with 1.0 mL of distilled water. After drying under a nitrogen gas stream, the lipid extract was dissolved in 1 mL chloroform for lipid analysis.

Following addition of internal standards (Devaiah et al., 2006), lipid species were quantified and identified by direct-infusion electrospray ionization tandem mass spectrometry (ESI–MS/MS) using a triple quadrupole instrument (API 4000, Applied Biosystems) at the Kansas Lipidomics Research Center, Kansas State University. Lipid classes were analyzed in class-specific ionization modes as follows: PC, LPC, sphingomyelin, PE, and LPE were detected as singly charged positive ions ([M+H]⁺); monogalactosyldiacylglycerol, digalactosyldiacylglycerol, phosphatidylglycerol, phosphatidylinositol, and phosphatidic acid were analyzed as [M+NH_4_]⁺ adducts; and phosphatidylserine and LPG were analyzed in negative ion mode as [M–H]⁻ ions. Additional lipid classes were analyzed using precursor ion or neutral loss scans, as previous described (Xiao et al., 2010).

### Antimicrobial activity of PLA2α

The mature form of rPLA2α was expressed in *E. coli* as a DsbC-fusion protein, purified by affinity chromatography followed by removal of the DsbC tag to obtain the free rPLA2α, as described previously (Ryu et al., 2005). To determine the antimicrobial activity of rPLA2α, the mature form (3 μg) produced as described (Ryu et al., 2005) was added to a suspension of virulent *Pst* (5 × 10^5^ CFU mL^-1^) in 100 μL of Tris–HCl (50 mM, pH 8.0) containing 10 mM CaCl_2_. Samples were incubated for 6 h at 28°C with gentle agitation, and surviving bacteria were titered by CFU assay.

### Release of lipid products from bacteria

To investigate whether PLA2α could generate lipid products such as LPE from bacterial membranes, bacterial lipid extracts (in 300 μL of Tris–HCl buffer, 50 mM, pH 8.0, with 10 mM CaCl_2_) were incubated with rPLA2α (3 μg per 100 μL) at 28°C for 15 min with gentle agitation. Lipids were then extracted as described previously (Lee et al., 2003). The mock control consisted of bacteria treated without rPLA2α. Lipid species were quantified by ESI–MS/MS as mentioned above.

In order to obtain direct evidence for the release of lipid products from invading bacteria, *Pst AvrRpm1* was cultured in medium containing [2-^14^C]-ethanolamine (10 μCi mL^-1^, 56 mCi mmol^-1^; Revvity, USA). Leaves of WT and *pla2α* plants were syringe-infiltrated with radiolabeled bacteria (1 × 10^7^ CFU mL^-1^ in H_2_O). Leaves were then harvested 3 h post-inoculation, and total lipids were extracted and separated by TLC (Ryu and Wang, 1996). Control (0 h) samples were harvested immediately after infiltration. Lipid zones on TLC corresponding to PE and LPE standards were collected, and radioactivity was measured using a liquid scintillation counter (PerkinElmer).

### Treatments with lipids

All phospholipids were purchased from Avanti Polar Lipids Inc. (Alabaster, AL, USA), and FFAs (16:0, 18:0, and 18:1) were obtained from Sigma-Aldrich. After solvent evaporation under nitrogen, lipids were resuspended by sonication in 0.018% Silwet L-77 to a final concentration of 100 μM. Unless otherwise stated, lipids were applied by spray throughout this study. Depending on the purpose, droplet or infiltration application was also used, as described below.

For gene expression assays in response to lipids, plant leaves were treated with lipids (100 μM in 0.018% Silwet L-77) either by spray inoculation or by applying four 10-μL droplets per leaf (equivalent to 4 nmol per leaf). For complementation of the macroHR phenotypic defect in *pla2α* plants, lysophospholipids (40 μM in H_2_O) were syringe-infiltrated into leaves at 1.5 h after syringe infiltration of *Pst AvrRpm1* (1 × 10^7^ CFU mL^-1^ in H_2_O). In all the other experiments, lipids or mock solutions were sprayed onto the leaf surfaces.

To complement impaired ETI-mediated resistance against *Pst AvrRpm1* in *pla2α* plants, lipids (LPE18:1, LPC16:0, or LPG16:0; 100 μM in 0.018% Silwet L-77) were sprayed 6 h prior to spray inoculation with *Pst AvrRpm1* (4 × 10^8^ CFU mL^-1^ in 0.018% Silwet L-77 and 7 mM MgCl_2_), followed by CFU assay at 4 dpi. To assess whether exogenous LPE alone is sufficient to recapitulate the resistance conferred by ETI against virulent *Pst*, LPE18:1 (100 μM in 0.018% Silwet L-77) or mock solution was sprayed onto leaves of WT and *pla2α* plants 6 h prior to spray inoculation with virulent *Pst* (1 × 10^7^ CFU mL^-1^ in 0.018% Silwet L-77 and 7 mM MgCl_2_), followed by CFU assay at 4 dpi. To further assess this resistance phenotype visually, LPE18:1 (100 μM in 0.018% Silwet L-77) or mock solution was sprayed onto leaves of WT plants 6 h prior to spray inoculation with virulent *Pst* (3 × 10^8^ CFU mL^-1^ in 0.018% Silwet L-77 and 7 mM MgCl_2_), and representative photographs were taken at 6 dpi.

### Treatments with rPLA2α protein and SA

To determine the lysophospholipid species generated by rPLA2α *in situ* and to evaluate its ability to induce *ICS1* and *PR1* expression in *pla2α* mutant leaves, rPLA2α protein (10 μg mL^-1^) was infiltrated at a dosage of 0.5 μg per leaf. Lipids and gene expression were analyzed at 3 and 6 hpi, respectively. Mock treatments were conducted using 50 mM Tris-HCl buffer (pH 8.0). For enzyme inactivation, rPLA2α was pre-incubated with 2 μM manoalide (an irreversible PLA2 inhibitor) at 37°C for 30 min, with inactivation confirmed via *in vitro* activity assay (Lee et al., 2010). For SA treatments, 4-week-old *Arabidopsis* plants were sprayed with 0.1 mM SA containing 0.018% Silwet L-77, while dH2O with 0.018% Silwet L-77 served as the mock control.

### Quantification of SA

Free SA was quantified as previously described (Bowling et al., 1994; Heck et al., 2003). Briefly, 0.5 g of fresh *Arabidopsis* leaf tissue was harvested after treatment with *Pst AvrRpm1* (1 × 10⁸ CFU mL^-1^ in 0.018% Silwet L-77) or LPE18:1 (100 μM in 0.018% Silwet L-77), extracted with methanol, and analyzed by HPLC with fluorescence detection. SA levels were quantified using external standards and normalized to fresh weight.

### MicroHR, macroHR, and ion leakage (cell death) assays

To assess microHR cell death induced by *Pst AvrRpm1*, *Arabidopsis* lines including Col-0 WT, *pla2α* and *pla2α-II*, and sid2, were spray-inoculated with *Pst AvrRpm1* (4 × 10^8^ CFU mL^-1^ in 0.018% Silwet L-77). After 12 h, leaves were collected, stained with trypan blue solution, destained in ethanol, and mounted in 60% glycerol (Fernández-Bautista et al., 2016). Images of cell death zones and whole leaves were captured using a stereo microscope (SZ61, Olympus, Japan). Full-leaf images were assembled by stitching multiple microscope fields of each section of the whole leaf using the “Stitch” function in microscope imaging software (eXcope, Dixi, South Korea).

For macroHR cell death assays, half of each leaf was syringe-infiltrated with a bacterial suspension (5 × 10^7^ CFU mL^-1^ in H_2_O). MacroHR symptoms were photographed at 15 hpi and 6 dpi. Mock-infiltrated leaves served as controls.

For the ion leakage assay with leaf discs, eight 6-mm discs were collected from fully expanded leaves of each *Arabidopsis* line (Col-0 WT, *pla2α,* and *pla2α-II*) and vacuum-infiltrated with either mock solution (10 mM MgCl_2_) or *Pst AvrRpm1* (5 × 10^7^ CFU mL^-1^ in 10 mM MgCl_2_) to induce synchronized ETI responses. After infiltration, discs were rinsed in water for 30 min and transferred to 6-well culture plate (SPL, South Korea) containing 6 mL ultrapure water, as described previously (Mackey et al., 2002; Ling et al., 2017). Electrolyte leakage was monitored at the indicated time points using a compact conductivity meter (LAQUAtwin-EC-11, Horiba, Japan).

For the ion leakage assay with intact leaves, half of each leaf was syringe-infiltrated with a bacterial suspension (5 × 10^7^ CFU mL^-1^ in H_2_O). At the designated time points, four leaves per sample were harvested, vacuum-infiltrated with distilled water for 5 min, and incubated at room temperature with gentle shaking for 1 h. Conductivity was measured using a conductivity meter (Mettler Toledo). Total electrolyte release was determined by freezing, followed by thawing to room temperature with shaking, and the data are expressed as percentages of this total leakage.

### Histochemical analysis of GUS activity

To visualize region-specific and NLR-dependent expression of *PLA2α* in *Arabidopsis* leaves following pathogen challenge, histochemical analysis of GUS activity was conducted as previously detailed (Jung et al., 2012). pPLA2α*:GUS* plants were syringe-infiltrated on the abaxial surface with bacterial pathogens at 3 h after the onset of the dark period and subsequently harvested for staining 1.5 hpi. The harvested leaves were immersed in GUS staining buffer to detect promoter-driven reporter activity. Stained tissues were then decolorized in 96% (v/v) ethanol to eliminate chlorophyll and subsequently transferred to 50% (v/v) glycerol for preservation and imaging.

### Gene expression analysis

*Arabidopsis* leaves were pre-flash-frozen in liquid nitrogen (6 leaves per sample) and homogenized using a TissueLyser II (Qiagen, Germany). Total RNA was then isolated using TRI Reagent® (Molecular Research Center, Inc. USA). Plant genomic DNA was removed using RQ1 DNase I (Promega, Korea). Real-time qPCR was performed using gene-specific primers (Table S4). Reference genes were *ACT1* or *PP2A* (At1g13320) (Hong et al., 2010).

### Transcriptomic and bioinformatic analyses

Total RNA was extracted from the leaves of 4-week-old *Arabidopsis* plants (Col-0 WT and *pla2α*) 12 h after spraying with one of the following solutions: Mock (dH_2_O containing 0.018% Silwet L-77, also used to prepare other solutions), 100 µM LPE18:1, *Pst AvrRpm1* suspension (1 × 10^8^ CFU mL^-1^) or 0.1 mM SA. RNA extraction was performed using TRI Reagent® (Molecular Research Center, Inc., USA) and plant genomic DNA was removed using the TURBO DNA-free™ Kit (Invitrogen, USA). Strand-specific cDNA libraries were prepared using the DNBSEQ eukaryotic mRNA library kit, and sequenced (PE150) on the DNBSEQ platform (BGI, Hong Kong, China). Sequencing yielded approximately 7.1 Gb of clean data per sample, which were filtered using SOAPnuke software with the following parameters: “-n 0.01 -l 20 -q 0.4 --adaML 0.25 --ada_trim --polyX 50 -minReadLen 150”. Raw reads were deposited in the Korea Sequence Read Archive (KRA) in Korea Bioinformation Center, Korea Research Institute of Bioscience and Biotechnology (accession KAP241514) at https://kbds.re.kr/KRA.

Cleaned reads were mapped to the *Arabidopsis* genome (TAIR10.5) and counted using CLC Genomics Workbench 10.1.1 with default settings (Qiagen, Germany). Differentially expressed genes (DEGs) were identified using DESeq2 on the iDEP2.01 platform with default settings (Ge et al., 2018). DEGs were selected based on adjusted *p*-value (adjPval) ≤0.05 and an absolute fold change ≥2. The enrichment of Gene Ontology (GO) and KEGG pathway, and MapMan categories was analyzed using DAVID (Sherman et al., 2022) and PlantGSAD (Ma et al., 2022), with thresholds of gene count ≥ 5 and a FDR ≤ 0.05. Enrichment figures were created in Origin 9.0 (Origin Lab, USA). Proportional Venn diagrams were drawn using DeepVenn (Hulsen, 2022). Genes responsible for the regulation and biosynthesis of SA, as well as SA downstream targets, were referenced from previous papers (Gonzalez-Bayon et al., 2019; Huang et al., 2020). Heatmap were visualized with GraphPad Prism 8 based on log_2_(fold change) (GraphPad Software Inc., USA).

### Calculation of gene expression levels from RNA-seq analysis

For selected genes of interest, expression levels were derived from processed RNA-seq data (log_2_ scale) using the iDEP2.01 platform with default settings. Log_2_(fold change) values were calculated by subtracting the mean log₂ normalized count of mock-treated replicates (n = 5) from that of each sample, and then converted to linear scale [fold change = 2^log_2_(FC)].

### ROS production assay

ROS production was assessed using two complementary assays: a luminol-based assay to monitor rapid ROS bursts in leaf discs (Smith and Heese, 2014; Yuan et al., 2021; Hong et al., 2023) and a ferric-xylenol orange (FOX) assay to quantify H_2_O_2_ accumulation in intact leaves (Bindschedler et al., 2006).

For the luminol-based assay, leaf discs (4 mm diameter, n = 10 per replicate) from WT and *pla2α* mutant plants were washed twice with sterile water and incubated overnight in a white 96-well plate (SPL, 30196) containing sterile water under gentle shaking and continuous light at 22°C. After an additional wash with sterile water, leaf discs were incubated in a reaction solution containing 20 μg mL^-1^ horseradish peroxidase (Sigma, P6782) and 200 μM luminol (Sigma, A8511), supplemented with a bacterial suspension of *Pst AvrRpm1* (5 × 10⁷ CFU mL^-1^). For the control group, sterile water (mock) was used instead of the bacterial suspension. Luminescence was measured using a VICTOR X3 multilabel plate reader with an integration time of 1 s per well and measurements taken at 2-min intervals over the measurement period. Relative luminescence units (RLU) were normalized as RLU_Pathogen − RLU_Mock, and total ROS production was calculated as the sum of RLUs over the measurement period. ROS production was analyzed in three phases: PTI (0–60 min), early ETI (ETI-I, 60–165 min), and late ETI (ETI-II, 165–390 min) phases.

For the FOX assay, three fully expanded leaves from WT and *pla2α* mutant plants infiltrated with mock, 100 μM LPE, or *Pst AvrRpm1* (1 × 10⁷ CFU mL^-1^) were harvested at the indicated time points and homogenized in ice-cold extraction buffer (50 mM potassium phosphate buffer, pH 6.5). Homogenates were centrifuged, and 100 μL of the supernatant was mixed with 900 μL of FOX reagent (250 μM ammonium ferrous sulfate, 100 μM xylenol orange, 100 mM sorbitol, and 25 mM H_2_SO_4_) and incubated for 30 min in the dark. H_2_O_2_ concentrations were determined at 560 nm using a standard curve generated from serial dilutions of H_2_O_2_.

### H_2_O_2_ floating assay

Fully expanded true leaves from 4-week-old WT and *pla2α* mutant plants were excised and placed abaxial side down in 10 mM H_2_O_2_ solution in Petri dishes sealed with Parafilm. Plates were incubated in the growth room for 5 days and photographed.

### Phylogenetic tree, orthologues of *Arabidopsis* PLA2α, and prediction of protein domain

The phylogenetic tree was obtained from Lifemap using the corresponding NCBI taxonomy IDs and edited with TreeGraph 2 (Stover and Muller, 2010; de Vienne, 2016). Representative species for each group or clade were selected as described in a previous study (Liu et al., 2022), with minor modifications. Most orthologs of *Arabidopsis* PLA2α were retrieved from the NCBI protein database, except for the ortholog in *Selaginella moellendorffii*, which was identified via BLAST analysis against the Phytozome database (https://phytozomenext.jgi.doe.gov/). Animal PLA2α orthologs were retrieved from SHOOT.bio using *Arabidopsis* PLA2α as the query sequence. Protein domains were predicted using the NCBI CD-search and visualized with the gene structure view (advanced) function in TBtools (Chen et al., 2020). Conserved motifs were illustrated using CLC Genomics Workbench 10. Signal peptides and cleavage sites were predicted with SignalP-5.0 (Almagro Armenteros et al., 2019).

## QUANTIFICATION AND STATISTICAL ANALYSIS

Data represent means ± SEM or SD as indicated in figure legends. Statistical significance was analyzed using either one-tailed or two-tailed Student’s *t*-test (\**P* < 0.05 and \*\**P* < 0.01), or by one-way ANOVA followed by Fisher’s LSD test (*P* < 0.05), where different letters indicate statistically significant differences.

