## SupplementaryTables for "Secreted phospholipase A2α generates a pathogen-derived lysophospholipid to signal local immunity"

Supplementary tables 1-4

**Table S1. Numbers of DEGs in different comparison groups, related to Figure 5**

| Comparisons | Up-regulated genes | Down-regulated genes | Note (Genotypes or treatments) |
|---|---|---|---|
| *Pst AvrRpm1* vs. Mock/WT | 1944 | 1443 | WT |
| LPE vs. Mock/WT | 744 | 279 | |
| SA vs. Mock/WT | 741 | 43 | |
| *Pst AvrRpm1* vs. Mock/*pla2α* | 972 | 104 | *pla2α* |
| LPE vs. Mock/*pla2α* | 884 | 168 | |
| SA vs. Mock/*pla2α* | 895 | 129 | |
| Mock/WT vs. *pla2α* | 8 | 4 | Mock |
| *Pst AvrRpm1*/WT vs. *pla2α* | 676 | 638 | *Pst AvrRpm1* |
| LPE/WT vs. *pla2α* | 66 | 99 | LPE |
| SA/WT vs. *pla2α* | 11 | 16 | SA |

**Table S2. Expression of selected genes based on RNA-seq data, related to Figure 5**

| Symbol | Gene ID | Fold change | | | | | | Adjust *P* value | | | | | |
|---|---|---|---|---|---|---|---|---|---|---|---|---|---|
| | | ***Pst Avr Rpm1* /WT** | **LPE /WT** | **SA /WT** | ***Pst Avr Rpm1 /pla2α*** | **LPE /*pla2α*** | **SA /*pla2α*** | ***Pst Avr Rpm1*/ WT** | **LPE /WT** | **SA /WT** | ***Pst Avr Rpm1 /pla2α*** | **LPE /*pla2α*** | **SA /*pla2α*** |
| *PLA2α* | AT2G06925 | 0.64 | 0.66 | 0.69 | 0.55 | 0.29 | 0.29 | 3.74E-03 | 1.88E-02 | 6.23E-02 | 5.89E-01 | 2.36E-01 | 2.83E-01 |
| *PLA2β* | AT2G19690 | 1.07 | 0.96 | 0.87 | 1.08 | 1.07 | 0.99 | 6.59E-01 | 8.51E-01 | 4.54E-01 | 7.27E-01 | 7.26E-01 | 9.77E-01 |
| *PLA2γ* | AT4G29460 | NA | NA | NA | NA | NA | NA | NA | NA | NA | NA | NA | NA |
| *PLA2δ* | AT4G29470 | NA | NA | NA | NA | NA | NA | NA | NA | NA | NA | NA | NA |
| *PR1* | AT2G14610 | 83.63 | 20.80 | 64.90 | 12.84 | 25.72 | 61.30 | 1.80E-29 | 4.76E-13 | 1.78E-25 | 1.40E-09 | 3.82E-15 | 1.63E-24 |
| *GST1* | AT1G02930 | 13.95 | 3.33 | 5.96 | 11.11 | 4.81 | 9.16 | 1.28E-13 | 3.67E-03 | 9.34E-06 | 2.42E-10 | 1.12E-04 | 5.36E-09 |
| *VSP1* | AT5G24780 | 0.26 | 2.13 | 0.20 | 1.28 | 2.18 | 0.23 | 1.46E-02 | 2.63E-01 | 1.72E-02 | 7.89E-01 | 2.52E-01 | 2.79E-02 |
| *JMT* | AT1G19640 | 1.61 | 1.73 | 1.22 | 1.77 | 1.48 | 1.19 | 6.68E-02 | 6.74E-02 | 6.38E-01 | 7.43E-02 | 2.13E-01 | 6.70E-01 |
| *PAL1* | AT2G37040 | 0.51 | 0.80 | 0.78 | 0.75 | 0.57 | 0.54 | 3.54E-03 | 4.60E-01 | 5.01E-01 | 4.01E-01 | 3.89E-02 | 3.29E-02 |
| *ICS1* | AT1G74710 | 7.70 | 2.01 | 3.70 | 3.17 | 2.06 | 3.68 | 6.10E-23 | 3.93E-03 | 6.09E-09 | 2.75E-07 | 3.05E-03 | 3.16E-09 |

a. NA, not applicable.

**Table S3. Signal peptide and cleavage sites predicted by SignalP-5.0, related to Figure S9**

| Species | Protein ID | Prediction | SP(Sec/SPI)[a] | OTHER | CS Position |
|---|---|---|---|---|---|
| *A. thaliana* | NP_565337.1 | SP(**Sec/SPI**) | 0.998427 | 0.001573 | CS pos: 20-21. VSA-LN. Pr: 0.8987 |
| *B. Napus* | KAH0873268.1 | SP(Sec/SPI) | 0.998218 | 0.001782 | CS pos: 22-23. VSA-LN. Pr: 0.9229 |
| *P. trichocarpa* | XP_002325228.2 | SP(Sec/SPI) | 0.987114 | 0.012886 | CS pos: 31-32. VQA-LN. Pr: 0.9209 |
| *G. max* | NP_001241376.1 | SP(Sec/SPI) | 0.934655 | 0.065345 | CS pos: 29-30. ACA-LN. Pr: 0.6057 |
| *M. truncatula* | XP_003620949.2 | SP(Sec/SPI) | 0.872431 | 0.127569 | CS pos: 29-30. VYA-LN. Pr: 0.7714 |
| *P. sativum* | XP_050904534.1 | SP(Sec/SPI) | 0.981498 | 0.018502 | CS pos: 28-29. AYA-LN. Pr: 0.9483 |
| *V. vinifera* | CBI40130.3 | SP(Sec/SPI) | 0.998735 | 0.001265 | CS pos: 20-21. TLA-LN. Pr: 0.8984 |
| *Z. Mays* | NP_001339274.1 | SP(Sec/SPI) | 0.996856 | 0.003144 | CS pos: 29-30. AAA-LN. Pr: 0.9339 |
| *T. aestivum* | XP_044368105.1 | SP(Sec/SPI) | 0.96343 | 0.03657 | CS pos: 33-34. SSA-LN. Pr: 0.7622 |
| *O. sativa* | Q9XG81 | SP(Sec/SPI) | 0.997771 | 0.002229 | CS pos: 25-26. SRG-LN. Pr: 0.8664 |
| *B. distachyon* | XP_003559514.1 | SP(Sec/SPI) | 0.980141 | 0.019859 | CS pos: 29-30. AAA-LD. Pr: 0.6189 |
| *A. comosus* | XP_020105054.1 | SP(Sec/SPI) | 0.999274 | 0.000726 | CS pos: 23-24. SRA-LN. Pr: 0.9337 |
| *A. trichopoda* | XP_006826876.1 | SP(Sec/SPI) | 0.937444 | 0.062556 | CS pos: 29-30. VMA-LQ. Pr: 0.7281 |
| *P. sitchensis* | ABR18340.1 | OTHER | 0.01464 | 0.98536 | NA |
| *C. japonica* | GLJ41712.1 | SP(Sec/SPI) | 0.582676 | 0.417324 | CS pos: 34-35. AQA-LN. Pr: 0.3420 |
| *C. richardii* | KAH7297221.1 | SP(Sec/SPI) | 0.53826 | 0.46174 | CS pos: 29-30. SVG-LV. Pr: 0.1652 |
| *A. capillus-veneris* | KAI5061270.1 | SP(Sec/SPI) | 0.787135 | 0.212865 | CS pos: 35-36. SCA-LI. Pr: 0.3345 |
| *S. moellendorffii* | XP_024534346.1 | OTHER | 0.000441 | 0.999559 | NA |
| *P. patens* | XP_024358604.1 | SP(Sec/SPI) | 0.969566 | 0.030434 | CS pos: 33-34. ALS-LV. Pr: 0.8163 |
| *M. polymorpha* | PTQ30484.1 | OTHER | 0.344715 | 0.655285 | NA |
| *E. debaryana* | KAG2485439.1 | SP(Sec/SPI) | 0.999888 | 0.000112 | CS pos: 21-22. AHA-ID. Pr: 0.9720 |
| *P. provasolii* | GHP12521.1 | SP(Sec/SPI) | 0.990966 | 0.009034 | CS pos: 24-25. ALA-GN. Pr: 0.6489 |
| *T. socialis* | PNH10697.1 | SP(Sec/SPI) | 0.999553 | 0.000447 | CS pos: 21-22. GKA-VD. Pr: 0.6928 |
| *V. carteri* | XP_002950584.1 | SP(Sec/SPI) | 0.999691 | 0.000309 | CS pos: 20-21. TGA-TD. Pr: 0.7522 |
| *C. reinhardtii* | XP_001699857.1 | SP(Sec/SPI) | 0.999768 | 0.000232 | CS pos: 24-25. VDA-VD. Pr: 0.7914 |
| *C. elegans* | O16654 | SP(Sec/SPI) | 0.978076 | 0.021924 | CS pos: 18-19. LNT-FV. Pr: 0.6062 |
| *D. rerio* | Q6DHQ7 | SP(Sec/SPI) | 0.655083 | 0.344917 | CS pos: 23-24. QRS-LR. Pr: 0.1008 |
| *M. musculus* | Q9QXX3 | SP(Sec/SPI) | 0.996524 | 0.003476 | CS pos: 17-18. GFS-EA. Pr: 0.5101 |
| *H. sapiens* | O15496 | SP(Sec/SPI) | 0.956227 | 0.043773 | CS pos: 31-32. GSG-EA. Pr: 0.4858 |

a. SP (Sec/SPI) indicates the probability that the sequence contains a "standard" secretory signal peptide, which is transported by the Sec translocon and cleaved by Signal Peptidase I (Lep). OTHER indicates the probability that the sequence does not contain any type of signal peptide. CS, predicted cleavage site; Pr, prediction probability; NA, not applicable.

**Table S4. Primers used in the real-time qPCR analysis and molecular cloning, related to method**

| Genes | Forward | Reverse | Product Size(bp) |
|---|---|---|---|
| **For real-time PCR** | | | |
| *ACT1* | CGTACTACCGGTATTGTGCTCGACT | GACAATTTCACGCTCTGCTGTGG | 189 |
| *PP2A* | GCGGTTGTGGAGAACATGATACG | GAACCAAACACAATTCGTTGCTG | 162 |
| *PLA*$_{2}$*α* | TCCATTTCCTTGACTAAAGAATG | AGATAATCATTATTCTTGGATTGG | 191 |
| *PLA*$_{2}$*β* | CGACGACGATGATGTTTCGC | CTCCTCACCAGGACAACCAG | 194 |
| *PLA*$_{2}$*γ* | GGATTCTCCACACAATGCCC | TGTCCACTTTACTCCGAGCG | 160 |
| *PLA*$_{2}$*δ* | CACGGAGTAAAGCGGACACA | CAAAACTTGAGAGTTATGAGTCACC | 135 |
| *PR1* | CATGTGGGTTAGCGAGAAGGCTA | CTCACTTTGGCACATCCGAGTCT | 120 |
| *ICS1* | CTAACCAGTCCGAAAGACGACCTC | CTTCCTTCGTAAGTCTCCCTGCC | 169 |
| *PAL1* | GAACTTATTAGATTCCTTAACGCCGG | GGAAACTGGTAATTGCTTCGAGAATC | 166 |
| *JMT* | GGCCAAAGAGGGTATCATCGAG | CCTCACTGATACTCCCACCTTCC | 167 |
| *VSP1* | CCTCGAATCGAACACCATCT | GGCACCGTGTCGAAGTTTAT | 135 |
| *RBOHD* | GACGATGAGTACGTGGAGATCA | GGAGGTGGTGTTGTTGAGGCT | 159 |
| *RBOHF* | CATCTATCGCTCCGATTTCGCT | CTCTCGTCGTTGATTTGTGACCA | 159 |
| *GOX3* | TTTAAGAGTCACCACTCATCAGAGAGAT | AAAGTATTCGATTATATAGTTGGATGGGA | 100 |
| *PRX33* | GGTCAGGTAATCCAGTGTTGC | GCTCTCCGGGGCTCAC | 89 |
| *PRX34* | TAGGGTCGGGTAAACCTGTG | GTTGCTCTCTCCGGTGGT | 97 |
| *PRX4* | GCGTTTAGGGCTATCGCAGAC | AAGCCTACCTTTGAACGTGAGG | 165 |
| *PRX25* | GCGACGGAAACGCTATTCTA | CTCTAAGACGACTCGCATACTTC | 93 |
| *PER58* | TACGAGCCCCGACTCATTTG | CGCTCCTGTCGAAGAGAACA | 100 |
| *PRX62* | TCGGACCACTGTGGCATCTCA | GAGTTAGGTCCCGATAAAAGCAC | 126 |
| *APX1* | TTTCCACCCTGGAAGAGAGGAC | TCACAACCCTTGGTAGCATCAGG | 75 |
| *CAT2* | AAGTATCCAACTCCGCCTGCTG | TGGATGAATCGTTCTTGCCTCTC | 131 |
| *ZAT12* | CACGGTGACTACGTTGAAGAAATC | CTCCAACTTGAGATTCAAATTGTC | 97 |
| *OXI1* | ACGACGCTAAATTGCTTGCT | CCGTGAAGAGACGGAAAGAG | 157 |
| **For molecular cloning** | | | |
| CP-*PLA2α* | GGGATCCGCCATATTGGAGAGAATGTC G | GGGATCCTCCTCGAGACCCTATTGCTG | 2112 |
| CP-noSP-*PLA2α*-1 | CGGGATCCGCCATATTGGAGAGAATG | GAGCTGAACACCGACGTTAAGCATAGTT GAACTGGAAAGCTC | 1199 |
| CP-noSP-*PLA2α*-2 | GAGCTTTCCAGTTCAACTATGCTTAACG TCGGTGTCAGCTC | AACTGCAGTCCTCGAGACCCTATTGCTG ATTC | 892 |
| *PLA2α*-OE | CGGGATCCAAAAATGGCGGCTCCGATC A | CCTCGCGATTAGGGTTTCTTGAGGACTTT GCC | 758 |
| Promoter-*PLA2α* | AAGCTTGCCATATTGGAGAGAATGTCG | GGATCCCATAGTTGAACTGGAAAGCTCG G | 1189 |
