## Supplementary Figures for "Secreted phospholipase A2α generates a pathogen-derived lysophospholipid to signal local immunity"

Supplementary figures 1-9

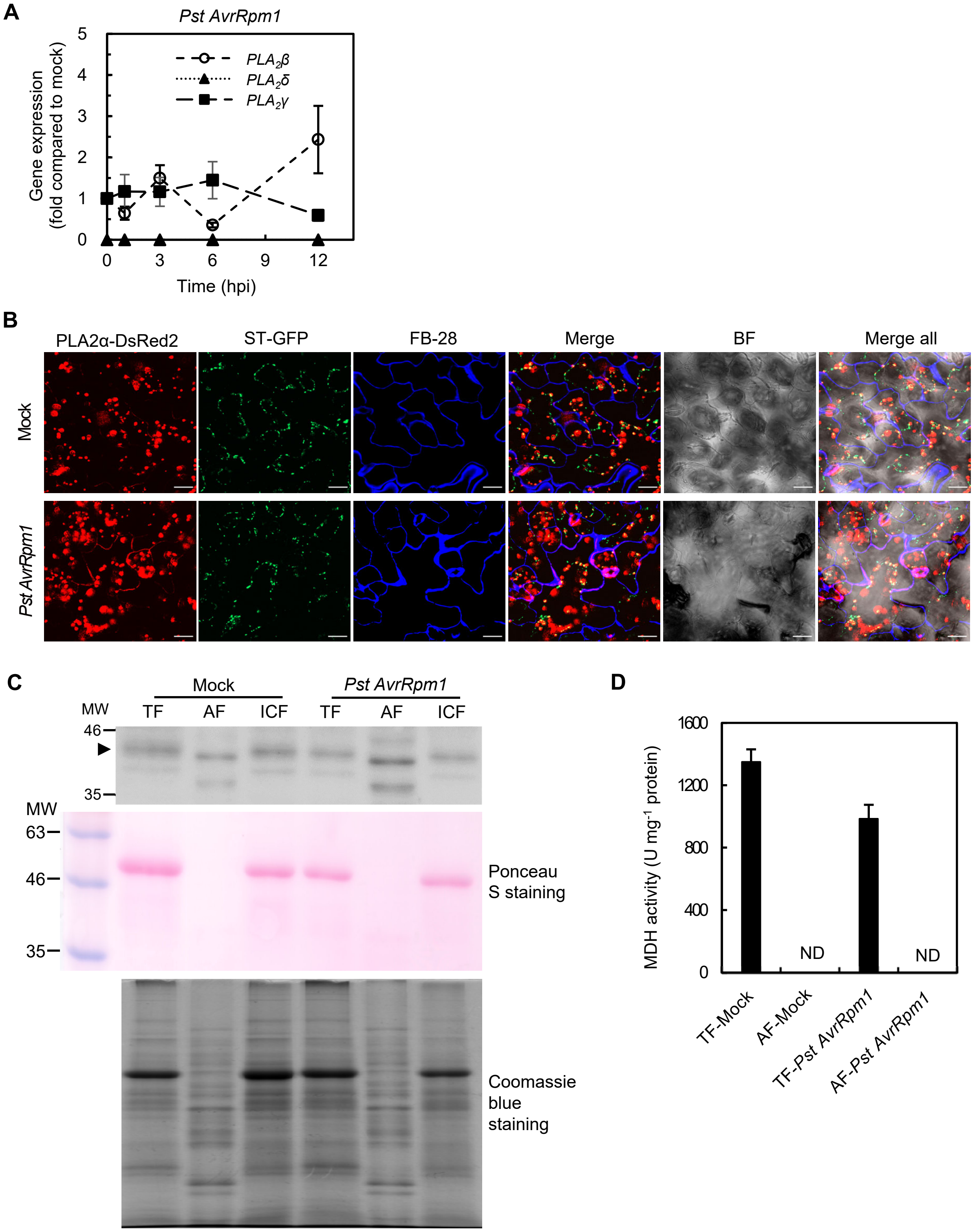


**Figure S1. Additional data, related to Figure 1**

**(A)** Time-course expression analysis of *PLA2β*, *PLA2γ*, and *PLA2δ* during ETI. Wild-type plants were spray-inoculated with *Pst AvrRpm1* ($1 \times 10^8$ CFU $mL^{-1}$), and gene expression in leaves was quantified by RT-qPCR at the indicated hpi. Data are expressed as fold-change relative to mock-inoculated plants.

**(B)** Full confocal image set corresponding to Figure 1B. Experiments were performed as described in Figure 1B. Individual fluorescence channels (PLA2α-DsRed2, red; ST-GFP, green; FB-28 cell wall stain, blue), merged overlay (Merge), bright-field (BF), and merged overlay with bright-field (Merge all) are shown for mock- (top row) and *Pst AvrRpm1*-inoculated (bottom row) leaves. "Merge" images are reproduced in Figure 1B. Scale bar, 20 μm.

**(C)** Loading controls for the immunoblot in Figure 1C. Experiments were performed as described in Figure 1C. The top panel is reproduced from Figure 1C. The middle panel shows Ponceau S staining of the blot membrane; the lower panel shows a parallel SDS-PAGE gel stained with Coomassie Brilliant Blue G-250. Gels were loaded with 3 μg of total fraction (TF) and intracellular fraction (ICF), and 1 μg of apoplastic fraction (AF).

**(D)** Malate dehydrogenase (MDH) activity in TF and AF as a cytoplasmic contamination control. Protein fractions used in (C) were assayed for MDH activity. ND, not detected.

Data represent means ± SD from at least three independent experiments.

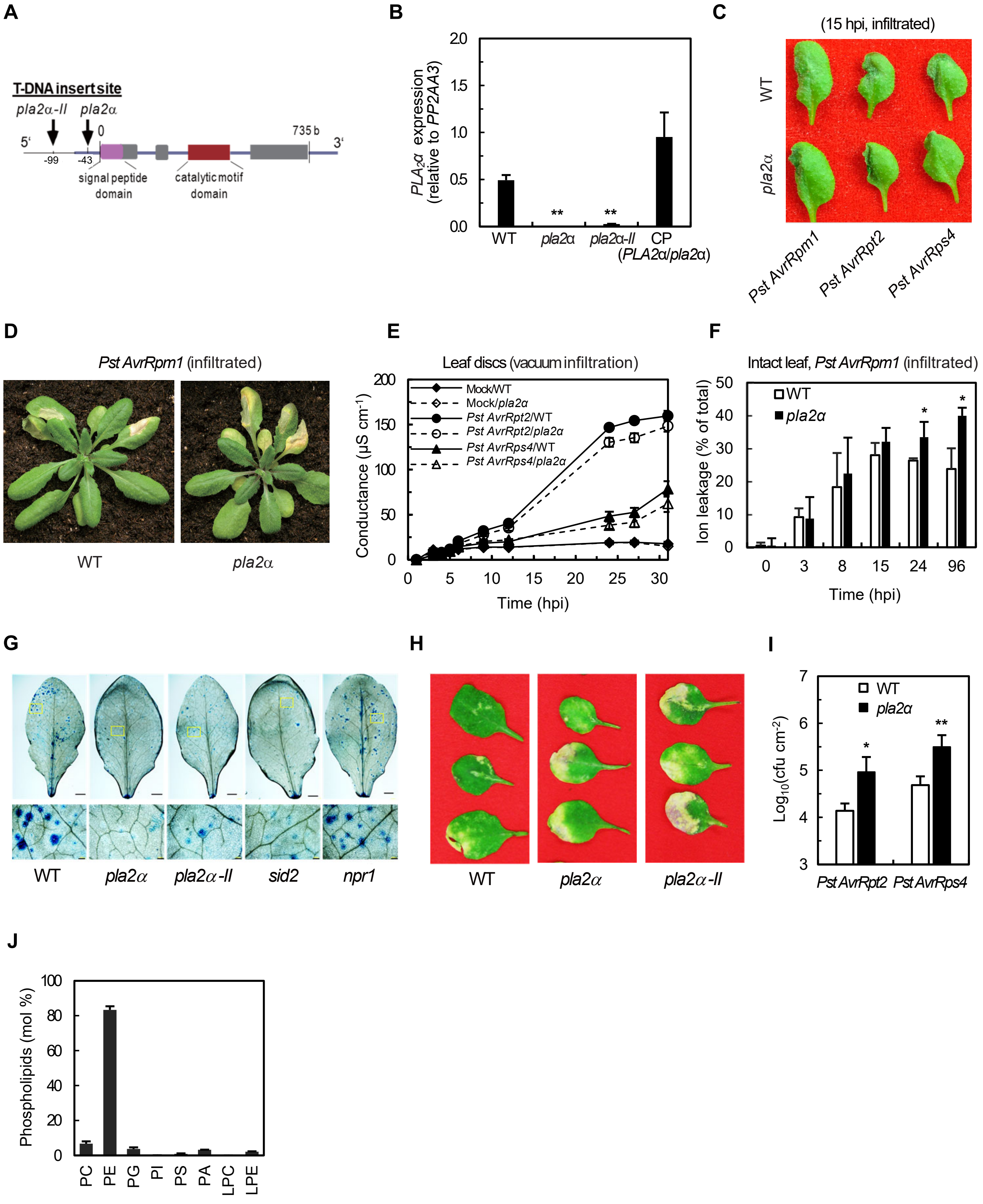


**Figure S2. Impaired disease resistance in *pla2α* mutants, related to Figure 1**

**(A)** Genomic structure of *PLA2α* (At2g06925) and T-DNA insertion sites in *pla2α* alleles. Exons, introns, and UTRs are indicated. Arrows mark T-DNA insertion positions in the 5′-UTR (*pla2α*; SALK_099415) and promoter region (*pla2α-II*; CS857021).

**(B)** *PLA2α* expression levels in WT, *pla2α*, *pla2α-II*, and complemented *pla2α* (CP). Leaf tissue expression was quantified by RT-qPCR and normalized to the reference gene *PP2AA3*. CP, *pla2α* complemented with *PLA2α* under its native promoter.

© MacroHR symptoms in intact leaves of WT and *pla2α* infiltrated with avirulent *Pst* strains (5 × $10^7$ CFU $mL^{-1}$) and photographed at 15 hpi.

**(D)** MacroHR symptoms in intact leaves of WT and *pla2α* syringe-infiltrated with *Pst AvrRpm1* (5 × $10^7$ CFU $mL^{-1}$) and photographed at 6 dpi.

**(E)** Time-course analysis of electrolyte leakage in WT and *pla2α* triggered by multiple avirulent *Pst* strains. Leaf discs were vacuum-infiltrated with *Pst AvrRpt2* or *Pst AvrRps4* ($5 \times 10^7$ CFU $mL^{-1}$) or mock solution, and conductance was monitored for up to ~30 hpi.
**(F)** Ion leakage in intact leaves of WT and *pla2α* during *Pst AvrRpm1* infection. Leaves were syringe-infiltrated with *Pst AvrRpm1* ($5 \times 10^7$ CFU $mL^{-1}$), and electrolyte leakage was measured from collected leaf tissue at the indicated hpi.
**(G)** MicroHR cell death in WT, *pla2α*, *sid2*, and *npr1*. Experiments were performed as described in Figure 1G. Lower panels show magnified images from the yellow-framed regions of the whole-leaf images above; lower panels are reproduced in Figure 1G. Scale bars: 2 mm (upper panels) and 200 μm (lower panels).
**(H)** Disease symptoms on rosette leaves of WT and *pla2α* at 6 dpi. Plants were spray-inoculated with *Pst AvrRpm1* ($4 \times 10^8$ CFU $mL^{-1}$) and photographed at 6 dpi.
**(I)** Bacterial growth in WT and *pla2α* inoculated with avirulent *Pst* strains. Plants were spray-inoculated with *Pst* AvrRpt2 or *Pst* AvrRps4 ($4 \times 10^8$ CFU $mL^{-1}$), and bacterial populations were quantified by CFU assay at 4 dpi.
**(J)** Phospholipid composition of *Pst AvrRpm1*. Total lipids were extracted from *Pst AvrRpm1* cultures and individual phospholipid classes were quantified by ESI-MS/MS.
Data represent means ± SD (E, and F) or ± SEM (B, I, and J) from at least three independent experiments. Asterisks indicate statistically significant differences by two-tailed Student's *t*-test (**P* < 0.05; ***P* < 0.01), except in (J), where a one-tailed Student's *t*-test was used.

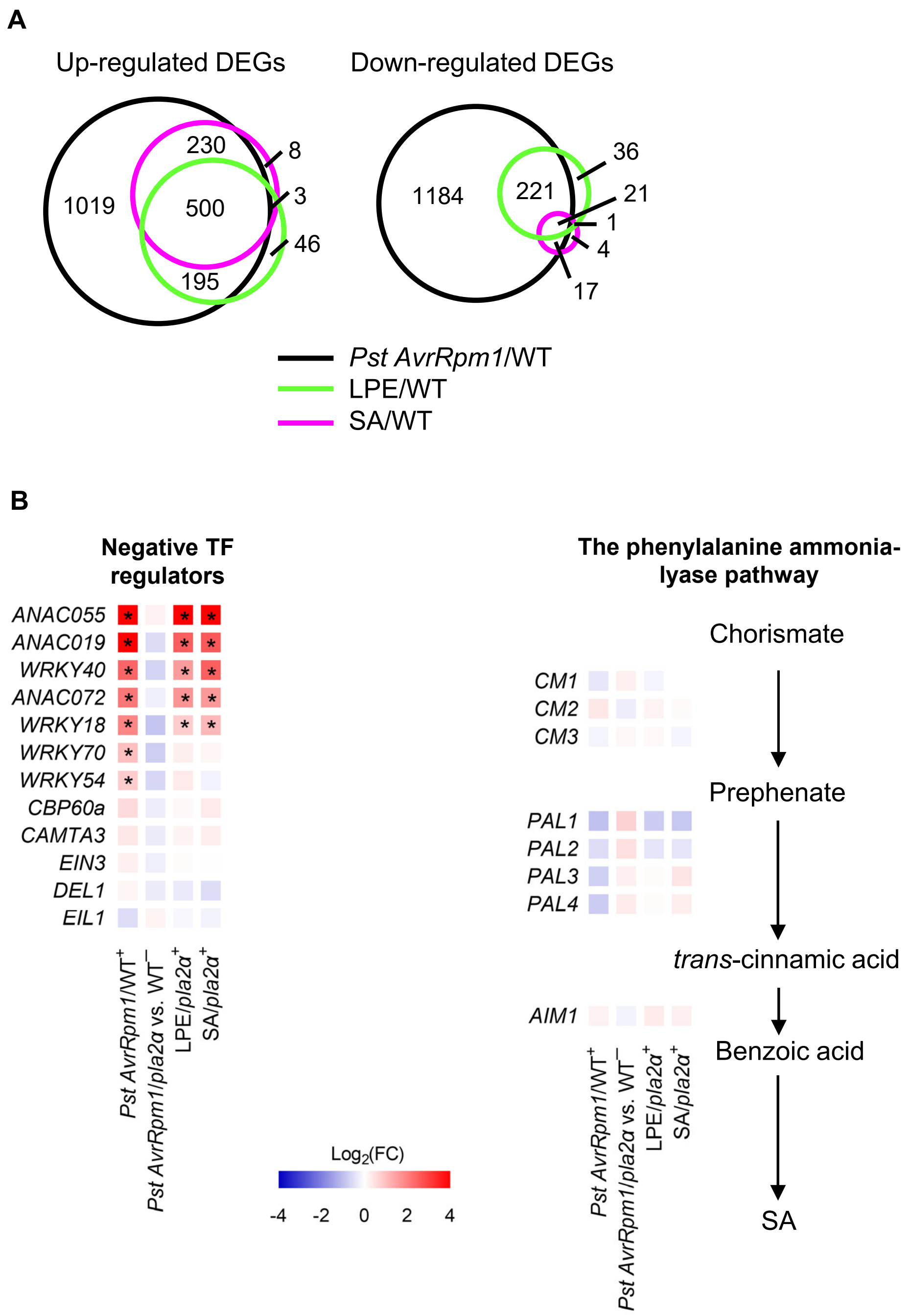


**Figure S3. Venn diagrams of DEGs and expression changes of genes involved in the SA signaling pathway across different treatment groups, related to Figure 5**

**(A)** Venn diagrams of DEGs in *Pst AvrRpm1*-, LPE18:1-, or SA-treated WT plants. RNA-seq analysis was performed as described in Figure 5A.

**(B)** Heatmaps of selected SA pathway genes across DEG sets from Figure 5. Left panel: transcription factors that negatively regulate SA biosynthesis, including NAC-domain proteins (*ANAC055*, *ANAC019*, *ANAC072*), WRKY transcription factors (*WRKY40*, *WRKY18*, *WRKY70*, *WRKY54*), and SA-signaling regulators (*CBP60a*, *CAMTA3*, *EIN3*, *DEL1*, *EIL1*). Right panels: genes of the ICS-independent SA biosynthesis branches, including chorismate mutase genes (*CM1*, *CM2*, *CM3*), phenylalanine ammonia-lyase genes (*PAL1–PAL4*), and the β-oxidation gene *AIM1*. Colors represent $\log_2$(fold change); asterisks indicate statistically significant DEGs. Upregulated DEGs are marked with "+" and downregulated DEGs with "−" in the column headers.

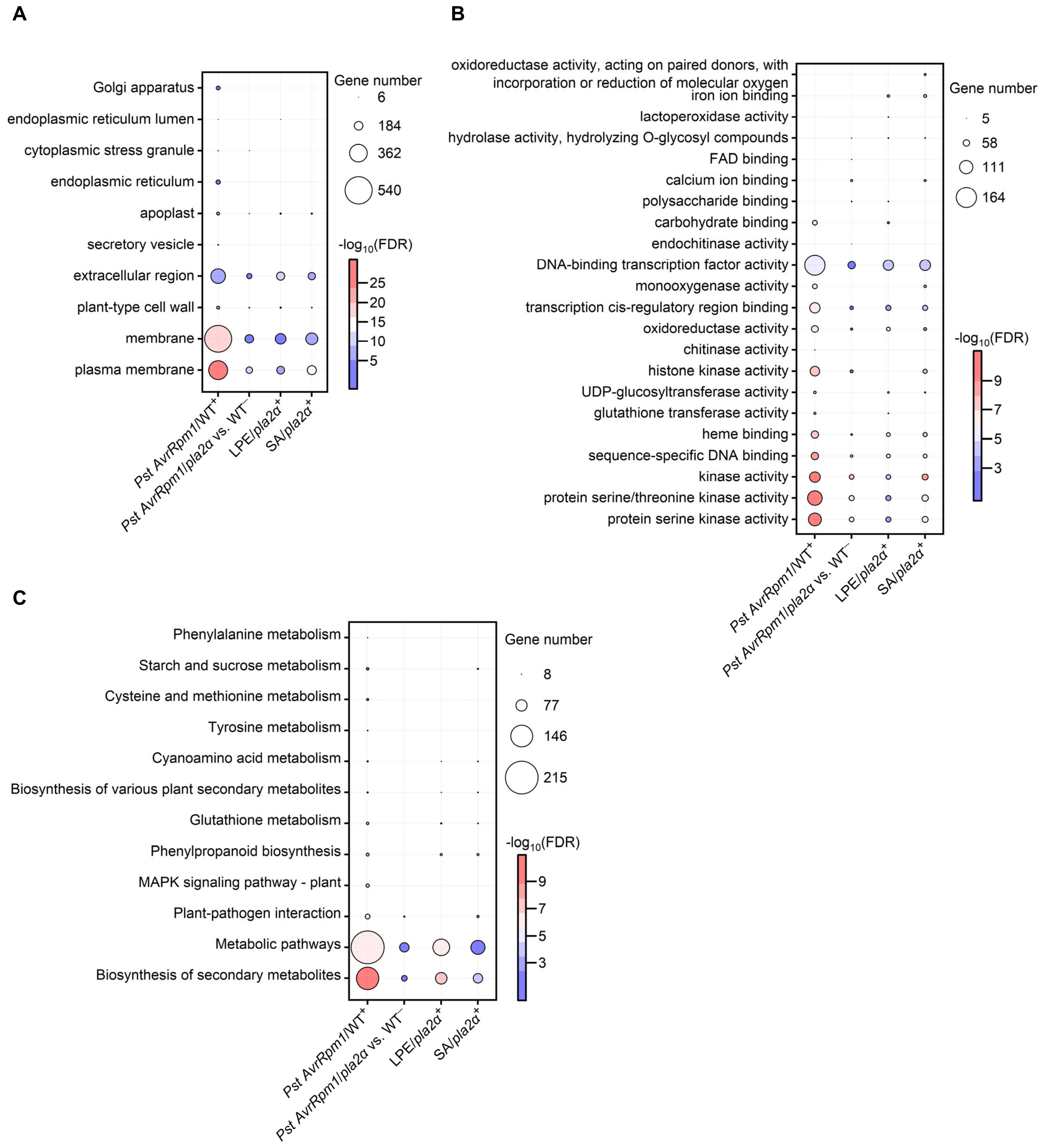


**Figure S4. GO term and KEGG pathway enrichment analysis of DEGs across treatment groups, related to Figure 5**

**(A–C)** Top 15 enriched GO terms and KEGG pathways among DEGs from the comparison groups shown in Figure 5C. GO cellular component terms (A), GO molecular function terms (B), and KEGG pathways (C) are shown. Bubble size represents the number of DEGs per term; color intensity represents statistical significance ($-\log_{10}$ FDR). Upregulated DEGs (+) were analyzed for all comparisons except *Pst AvrRpm1*/*pla2α* vs. WT$^-$, for which downregulated DEGs (−) were used. RNA-seq analysis was performed as described in Figure 5. Terms were included based on FDR ≤ 0.05 with a minimum of 5 genes per term.

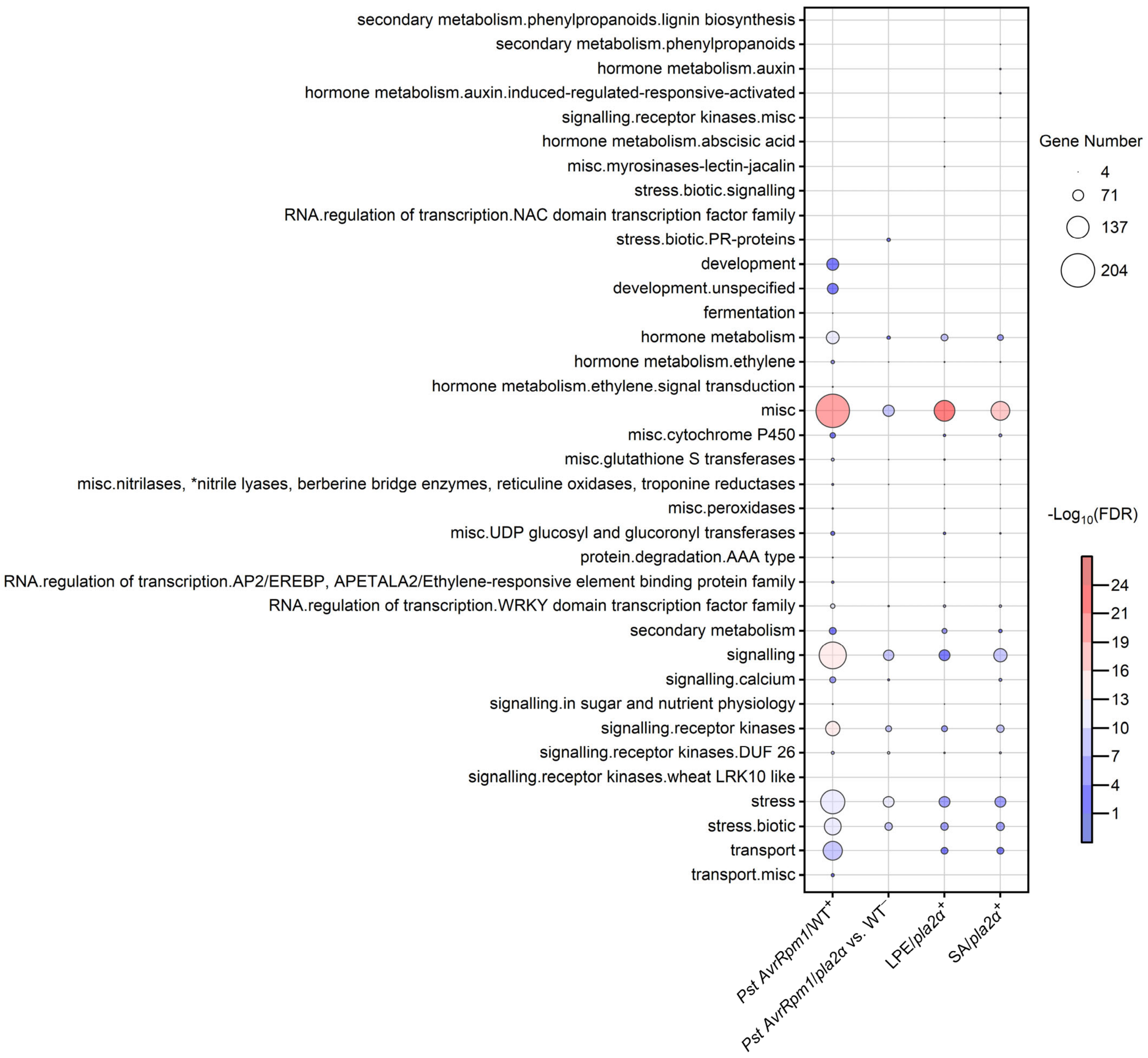


**Figure S5. MapMan functional category enrichment among DEGs across treatment groups, related to Figure 5**

Bubble plot showing enriched MapMan functional categories among DEGs from the comparison groups in Figure 5. Bubble size represents the number of DEGs per category; color intensity represents statistical significance ($-\log_{10}$ FDR). Upregulated DEGs (+) were analyzed for all comparisons except *Pst AvrRpm1*/*pla2α* vs. WT$^{-}$, for which downregulated DEGs (−) were used. Categories were included based on FDR ≤ 0.05 with a minimum of 5 genes per category. RNA-seq analysis was performed as described in Figure 5.

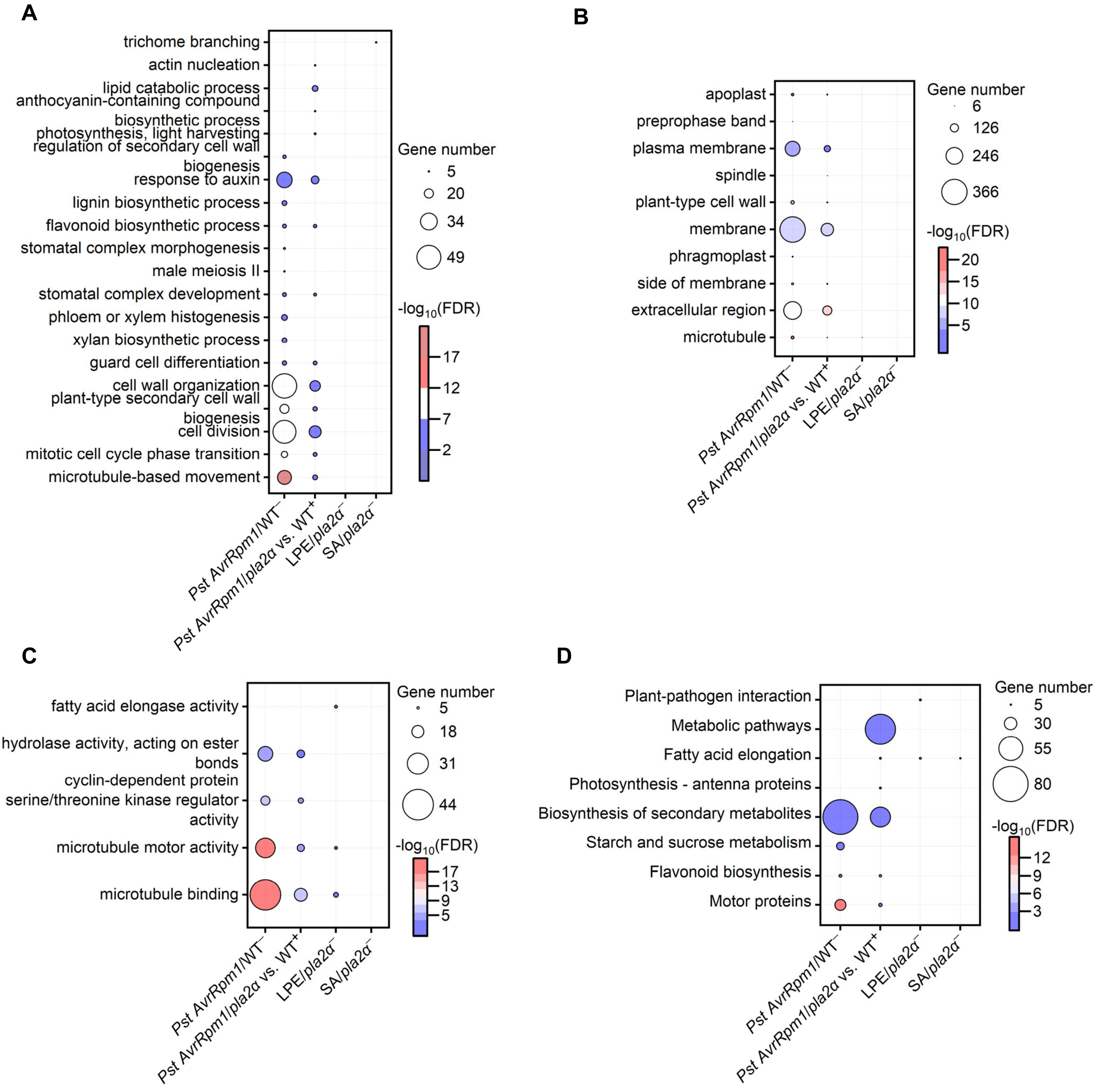


**Figure S6. Top 15 enriched GO terms and KEGG pathways among DEGs across treatment groups, related to Figure 5**

**(A–D)** Top 15 enriched GO terms and KEGG pathways among downregulated DEGs from the comparison groups shown in Figure 5. GO biological process terms (A), GO cellular component terms (B), GO molecular function terms (C), and KEGG pathways (D) are shown. Bubble size represents the number of DEGs per term; color intensity represents statistical significance ($-\log_{10}$ FDR). Downregulated DEGs (−) were analyzed for all comparisons except *Pst AvrRpm1*/*pla2α* vs. WT$^+$, for which upregulated DEGs (+) were used. Terms were included based on FDR ≤ 0.05 with a minimum of 5 genes per term. RNA-seq analysis was performed as described in Figure 5.

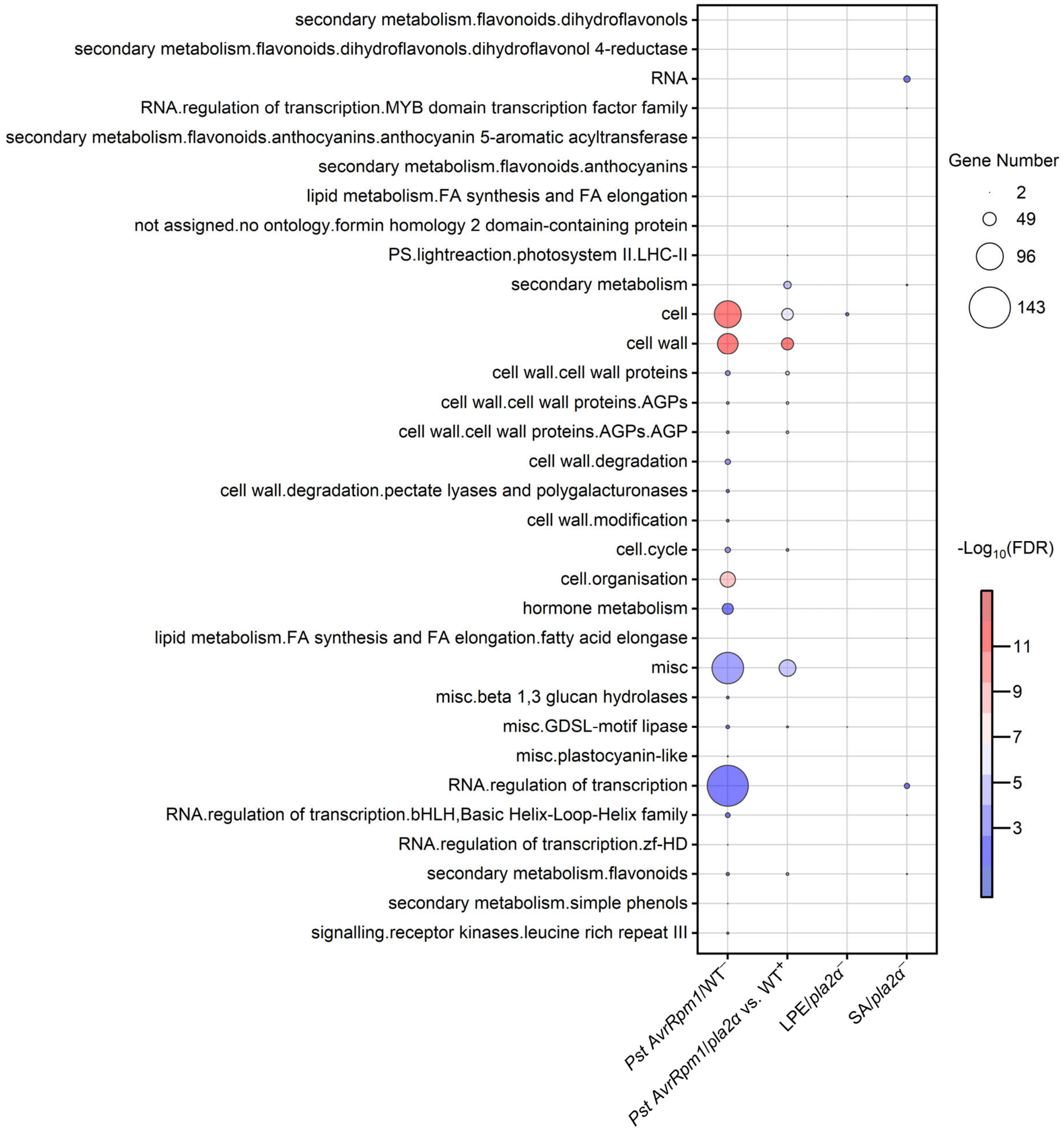


**Figure S7. MapMan functional category enrichment among downregulated DEGs across treatment groups, related to Figure 5**

Bubble plot showing enriched MapMan functional categories among downregulated DEGs from the comparison groups in Figure 5. Bubble size represents the number of DEGs per category; color intensity represents statistical significance ($-\log_{10}$ FDR). Downregulated DEGs (−) were analyzed for all comparisons except *Pst AvrRpm1*/*pla2α* vs. $WT^{+}$, for which upregulated DEGs (+) were used. Categories were included based on FDR ≤ 0.05 with a minimum of 5 genes per category. RNA-seq analysis was performed as described in Figure 5.

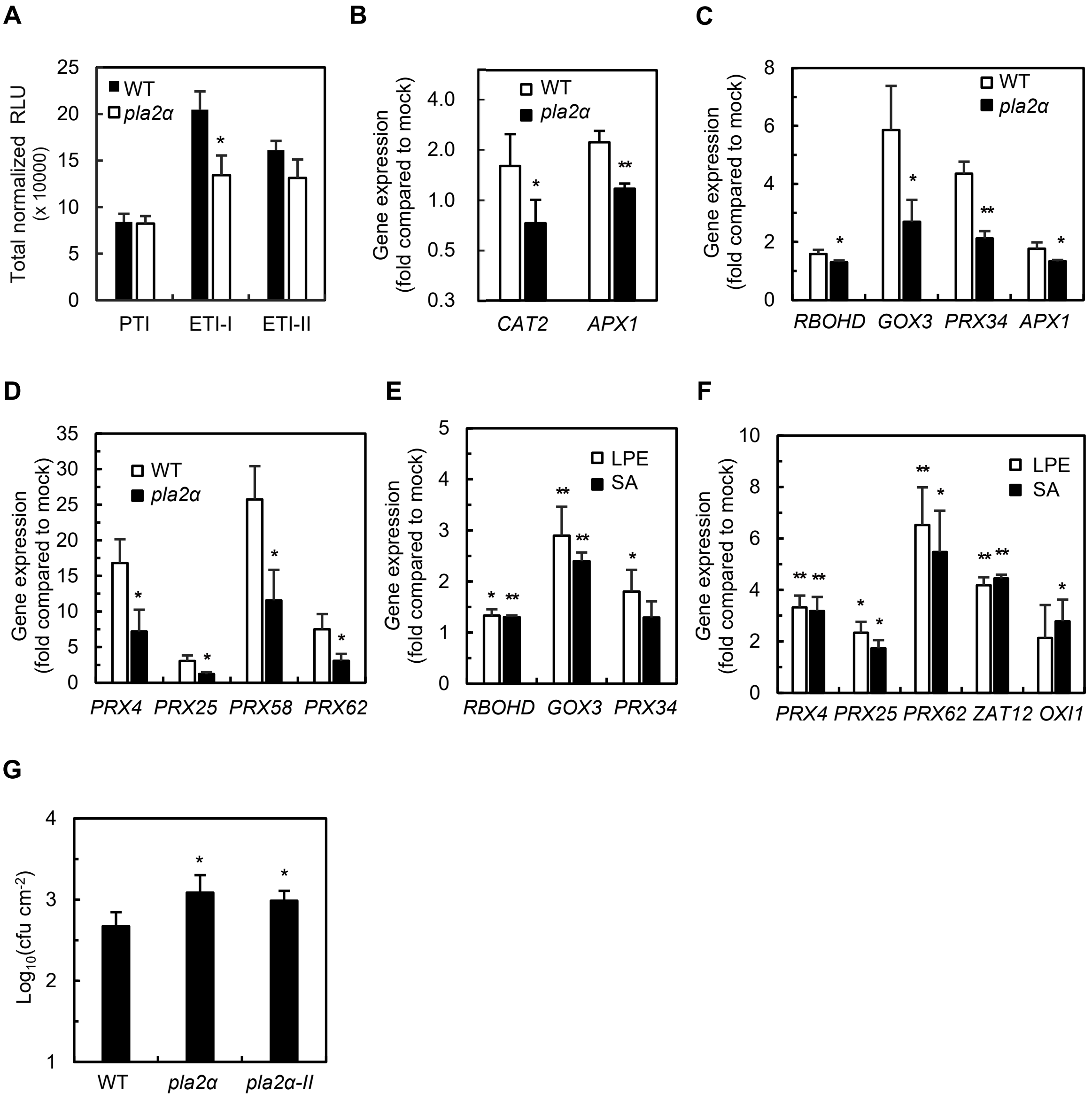


**Figure S8. Reduced $H_2O_2$ accumulation and altered ROS-associated gene expression in *pla2α*, related to Figure 6**

**(A)** Quantification of total ROS production during PTI and ETI phases in WT and *pla2α*. Data represent the area under the curves in Figure 6E, integrated over three intervals: PTI (0–60 min), ETI-I (60–165 min), and ETI-II (165–390 min).

**(B)** Expression of $H_2O_2$-scavenging genes in WT and *pla2α* leaf discs during ETI. Leaf discs were treated as in Figure 6E, and *CAT2* and *APX1* expression was quantified by RT-qPCR. The Y-axis is $\log_2$-transformed. Data are expressed as fold-change relative to mock-treated plants.

**(C and D)** Expression of ROS-producing genes in WT and *pla2α* during ETI, derived from RNA-seq data. Shown are *RBOHD*, *GOX3*, *PRX34*, and *APX1* (C) and *PRX4*/25/58/62 (D). Data are expressed as fold-change relative to mock-treated plants.

**(E and F)** Expression of ROS-related genes in LPE18:1- or SA-treated *pla2α*, derived from RNA-seq data. Shown are *RBOHD*, *GOX3*, and *PRX34* (E) and *PRX4/25/62*, *ZAT12*, and *OXI1* (F). Data are expressed as fold-change relative to mock-treated plants.

**(G)** Bacterial growth in the distant area of WT, *pla2α,* and *pla2α-II* leaves. Leaf tissue from the distant area as defined in Figure 6K was collected from leaves syringe-infiltrated with *Pst AvrRpm1* ($4 \times 10^8$ CFU $mL^{-1}$) and bacterial populations were quantified by CFU assay at 4 dpi.

Data represent means ± SD (G) or ± SEM (A–F) from at least three independent experiments. Statistical significance was assessed by two-tailed Student's *t*-test (A, B, and G) or one-tailed Student's *t*-test (C–F) ($^*P < 0.05$; $^{**}P < 0.01$).

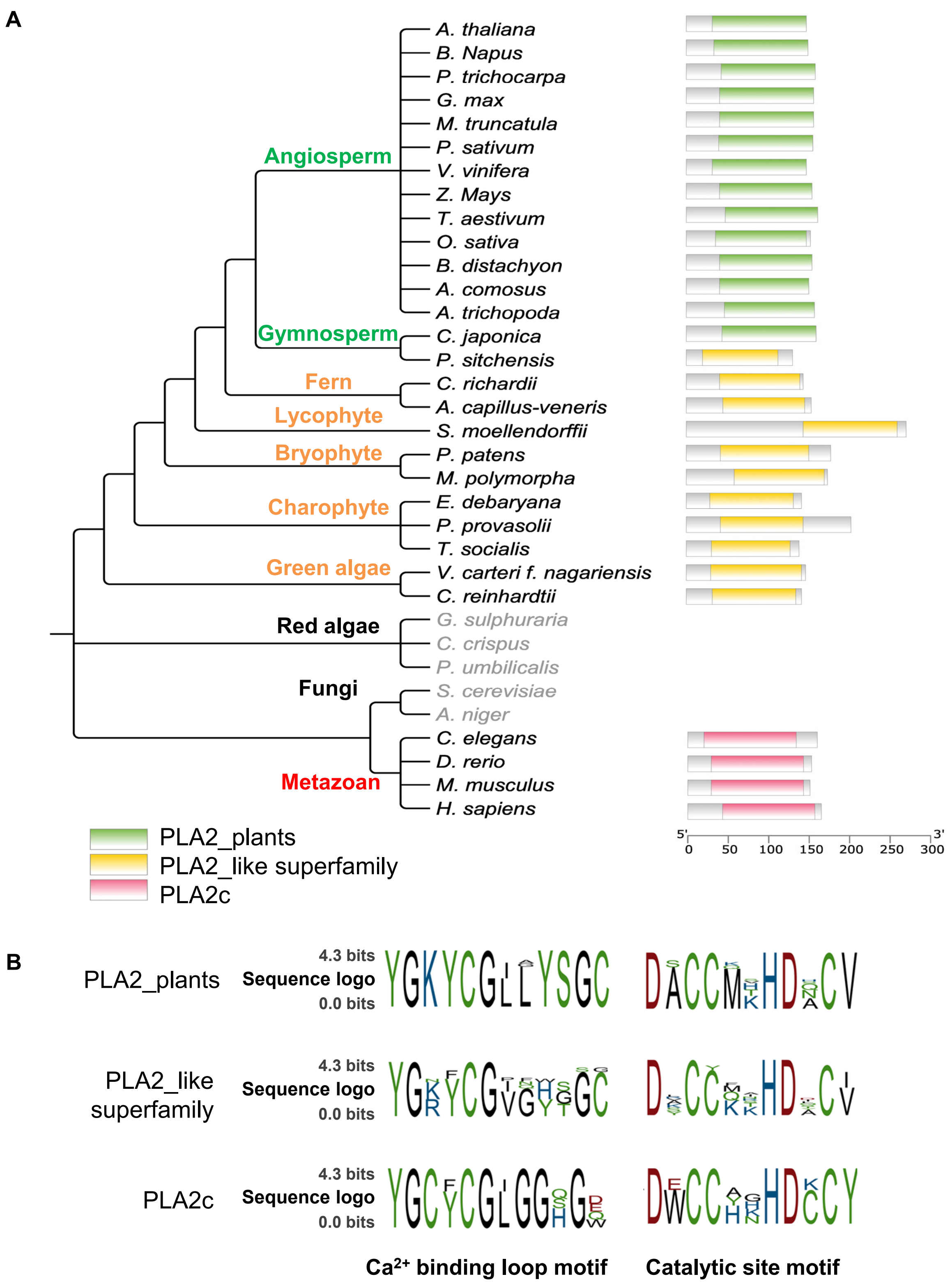


**Figure S9. Phylogenetic distribution and domain architecture of PLA2α orthologs across eukaryotes**
**(A)** Phylogenetic tree of PLA2α orthologs from representative eukaryotes. Major lineages are color-coded (seed-bearing plants, green; non-seed plants/algae, yellow; metazoans, red). Conserved domain annotations reveal lineage-specific architectures: PLA2_plants predominates in seed-bearing plants, while PLA2_like and PLA2c are characteristic of non-seed plant/algal and metazoan lineages, respectively.
**(B)** Sequence logos of conserved motifs within each PLA2 domain class. Logos were generated from aligned orthologs for PLA2_plants, PLA2_like, and PLA2c domains, with stack height reflecting the degree of sequence conservation at each position (bits).
See also Table S3.
